# Novel tumorigenic FOXM1-PTAFR-PTAF axis revealed by multi-omic profiling in *TP53/CDKN2A-*double knockout human gastroesophageal junction organoid model

**DOI:** 10.1101/2022.05.10.491356

**Authors:** Hua Zhao, Yulan Cheng, Andrew Kalra, Ke Ma, Yueyuan Zheng, Benjamin Ziman, Caitlin Tressler, Kristine Glunde, Eun Ji Shin, Saowanee Ngamruengphong, Mouen Khashab, Vikesh Singh, Robert A. Anders, Simran Jit, Nicolas Wyhs, Wei Chen, Xu Li, De-Chen Lin, Stephen J. Meltzer

## Abstract

Inactivation of the tumor suppressor genes *TP53* and *CDKN2A* occurs early during gastroesophageal junction (GEJ) tumorigenesis. However, due to a paucity of GEJ-specific disease models, cancer-promoting consequences of *TP53* and *CDKN2A* inactivation at the GEJ have been incompletely characterized. Here we report the development of the first wild-type primary human GEJ organoid model, as well as a CRISPR-edited transformed GEJ organoid model. CRISPR/Cas9 engineering to inactivate *TP53* and *CDKN2A* (*TP53/CDKN2A*^KO^) in GEJ organoids induced morphologic dysplasia as well as pro-neoplastic features *in vitro* and tumor formation *in vivo.* Notably, lipidomic profiling identified several Platelet-Activating Factors (PTAFs) among the most upregulated lipids in CRISPR-edited organoids; and importantly, PTAF/PTAFR abrogation by siRNA knockdown or a pharmacologic inhibitor (WEB2086) significantly blocked proliferation and other pro-neoplastic features of *TP53/CDKN2A*^KO^ GEJ organoids *in vitro* and tumor formation *in vivo*. In addition, murine xenografts derived from Eso26, an established esophageal adenocarcinoma (EAC) cell line, were suppressed by WEB2086. Mechanistically, *TP53/CDKN2A* dual inactivation disrupted both the transcriptome and the DNA methylome, likely mediated by key transcription factors, particularly Forkhead Box M1 (FOXM1). Importantly, FOXM1 activated PTAFR transcription by binding to the PTAFR promoter, further amplifying the PTAF-PTAFR pathway. In summary, we established a robust model system for investigating early GEJ neoplastic events, identified crucial metabolic and epigenomic changes occurring during GEJ model tumorigenesis, and discovered a potential cancer-therapeutic strategy, while providing insights into pro-neoplastic mechanisms associated with *TP53/CDKN2A* inactivation in early GEJ neoplasia.

**One Sentence Summary:** Novel tumorigenic FOXM1-PTAFR-PTAF axis revealed by multi-omic profiling in *TP53/CDKN2A-*double knockout human gastroesophageal junction organoid model.

**Graphic Abstract:** 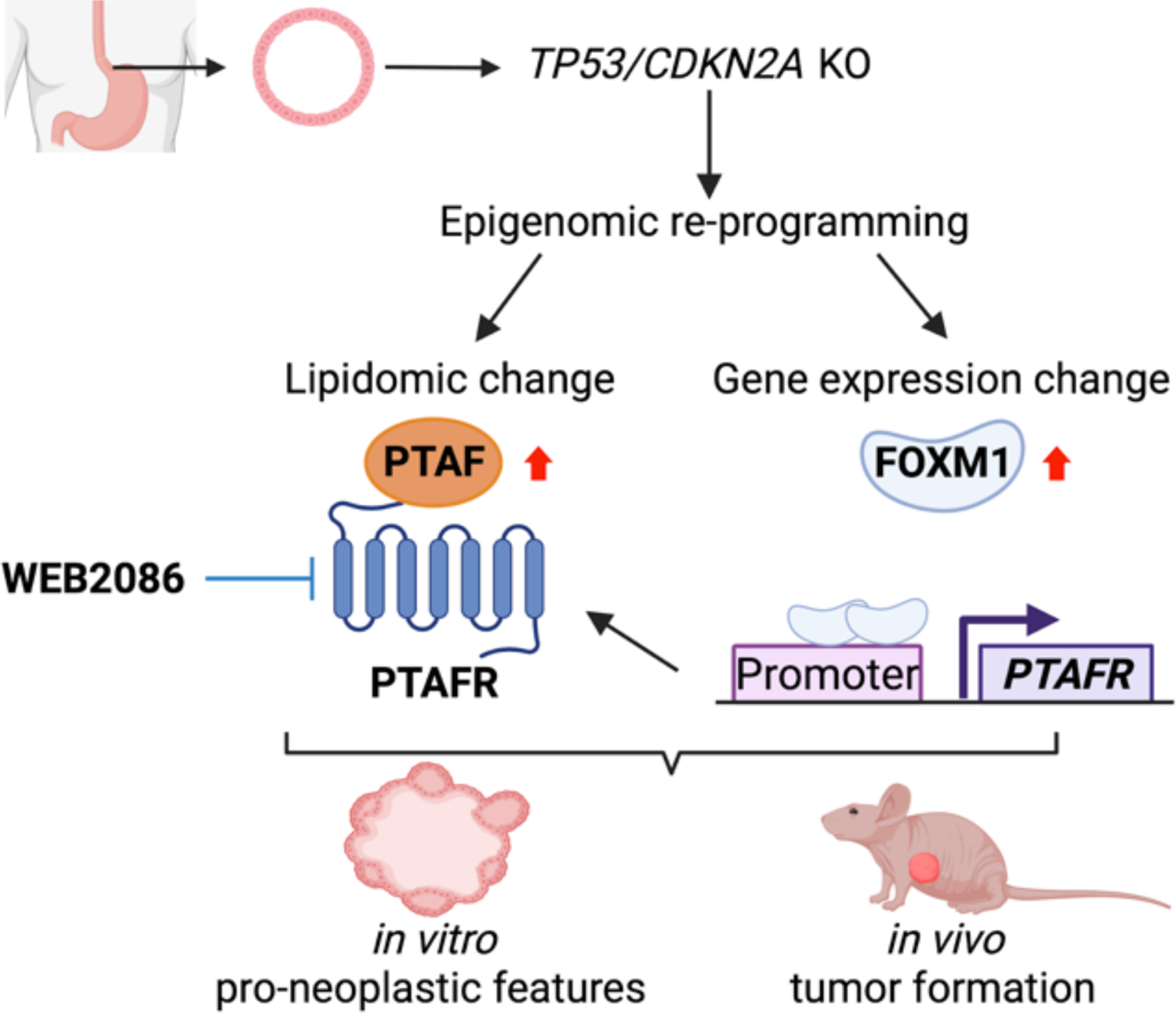

## INTRODUCTION

Gastroesophageal cancers account for over 1 million deaths annually, representing the second-highest cause of cancer death worldwide. One category of these neoplasms, gastroesophageal junction (GEJ) adenocarcinoma, has increased more than 2.5-fold in incidence in the United States and other Western countries in recent decades (*1, 2*). Compared with other gastroesophageal region cancers, GEJ tumors are particularly aggressive and have a dire prognosis; thus, vigorous investigation of their molecular basis is crucially needed. However, this goal has been difficult to achieve, in part due to a lack of biologically relevant GEJ-specific early disease models. Human organoids are robust models that recapitulate and maintain essential genetic, functional and phenotypic characteristics of their tissues of origin (*3*). Filling a knowledge gap between transgenic animal models and cell lines, organoids provide a valuable platform for understanding fundamental oncogenic mechanisms as well as identifying candidate cancer-therapeutic targets (*4*).

Lipid metabolic reprogramming is a known hallmark of cancer (*5, 6*), and certain lipid profiles have been proposed as either biomarkers for early cancer detection or targets for cancer treatment (*7–9*). Aberrations in the fate and composition of the lipidome promote tumorigenesis, enhance cancer cell growth, and support cancer cell survival in challenging microenvironments by activating oncogenic pathways and processes (*9*). Indeed, by performing lipidomic profiling in gastroesophageal cancers, we recently demonstrated that heightened synthesis of specific lipid species was required for cancer cell viability and proliferation (*10, 11*). However, metabolic changes in the lipidome of GEJ cancers are still largely unknown, while their biological significance awaits further investigation.

In the present study, we investigated GEJ tumorigenesis by establishing the first human normal GEJ-derived organoids, modified by *TP53/CDKN2A* dual-knockout (*TP53/CDKN2A*^KO^) using CRISPR/Cas9 genome editing. Utilizing this GEJ-specific disease model, we addressed the phenotypic, metabolic and epigenomic alterations during the early neoplastic transformation of GEJ cells, with the goal of providing insights into the mechanistic basis of this aggressive and poorly understood malignancy.

## RESULTS

### Establishment and characterization of human normal GEJ organoids

To address the paucity of biologically relevant GEJ-specific disease models, we generated three-dimensional (3D) organoids from human primary endoscopic GEJ biopsies, which were confirmed pathologically to contain neither dysplastic nor neoplastic cells. Briefly, freshly isolated GEJ crypts (prepared as described under *Methods*) were embedded in Matrigel and incubated with conditioned medium containing stem-critical growth factors (**Fig. 1A** and **Table 1**). Under culture conditions we established, 3D organoids were generated from 6 normal GEJ biopsies, with a 100% success rate. We characterized these organoids using phase-contrast imaging, hematoxylin/eosin (H&E) staining, and viability (WST-1) assays (**Fig. 1B-D**). At day 4 after initial seeding, 3D spherical structures were formed and reached 25 µm in diameter. These structures continued to grow, eventually reaching a size plateau of 106 µm between day 24 and 29 (**Fig. 1B and C**). These GEJ organoids consisted of 120-250 cells at day 24, indicating that population doubling time in culture was 77.85 ± 5.54 hrs. Histologic analyses revealed that normal GEJ organoids formed a 3D structure consisting of single-layered epithelial cells. The cell viability of organoids increased to a peak on day 14 and significantly diminished by day 24 (Fig 1D). We continued to monitor organoids from these 6 independent normal GEJ biopsies; they propagated continuously for 4 to 6 months *ex vivo*.

**Fig. 1.**
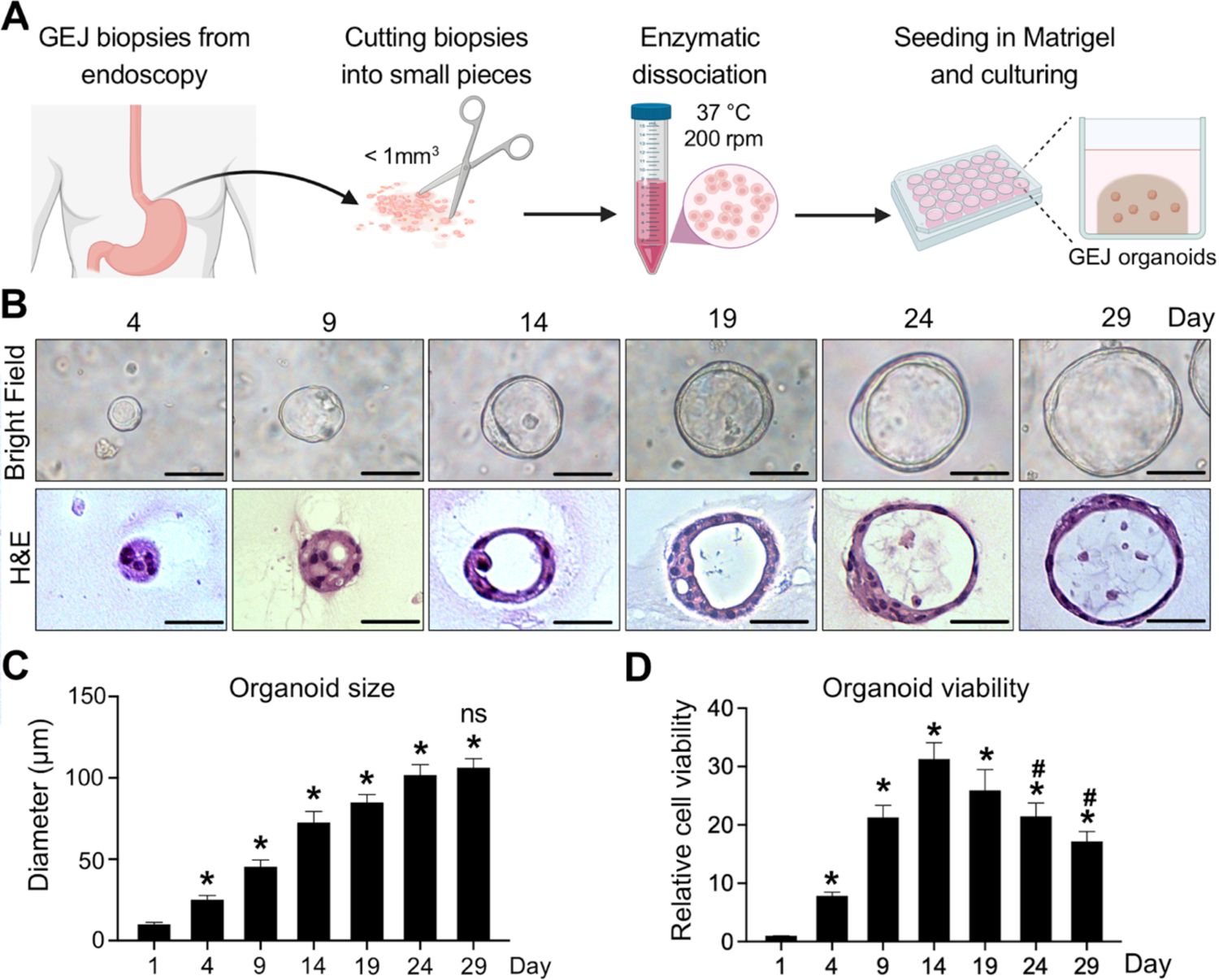
Establishment and characterization of human normal GEJ organoids. (A) A workflow of organoid generation from human primary endoscopic GEJ biopsies. Biopsies of normal GEJ mucosa were taken by upper endoscopy, and then minced and enzymatically dissociated. The cell suspension was mixed with Matrigel to initiate 3D organoid culture in the conditioned medium. (B-D) GEJ organoids were analyzed for structural and growth properties at the indicated time points. 3D organoids were photomicrographed under phase-contrast microscopy (B, upper panel) and collected for H&E staining (B, lower panel). Scale bar = 50 µm. Average organoid size (C) and viability (D) were determined at each timepoint. *, P < 0.05 *vs*. Day 1; #, P < 0.05 *vs.* Day 14; ns, not significant *vs.* Day 24. Data represent 6 biological replicates.

**Table 1.**
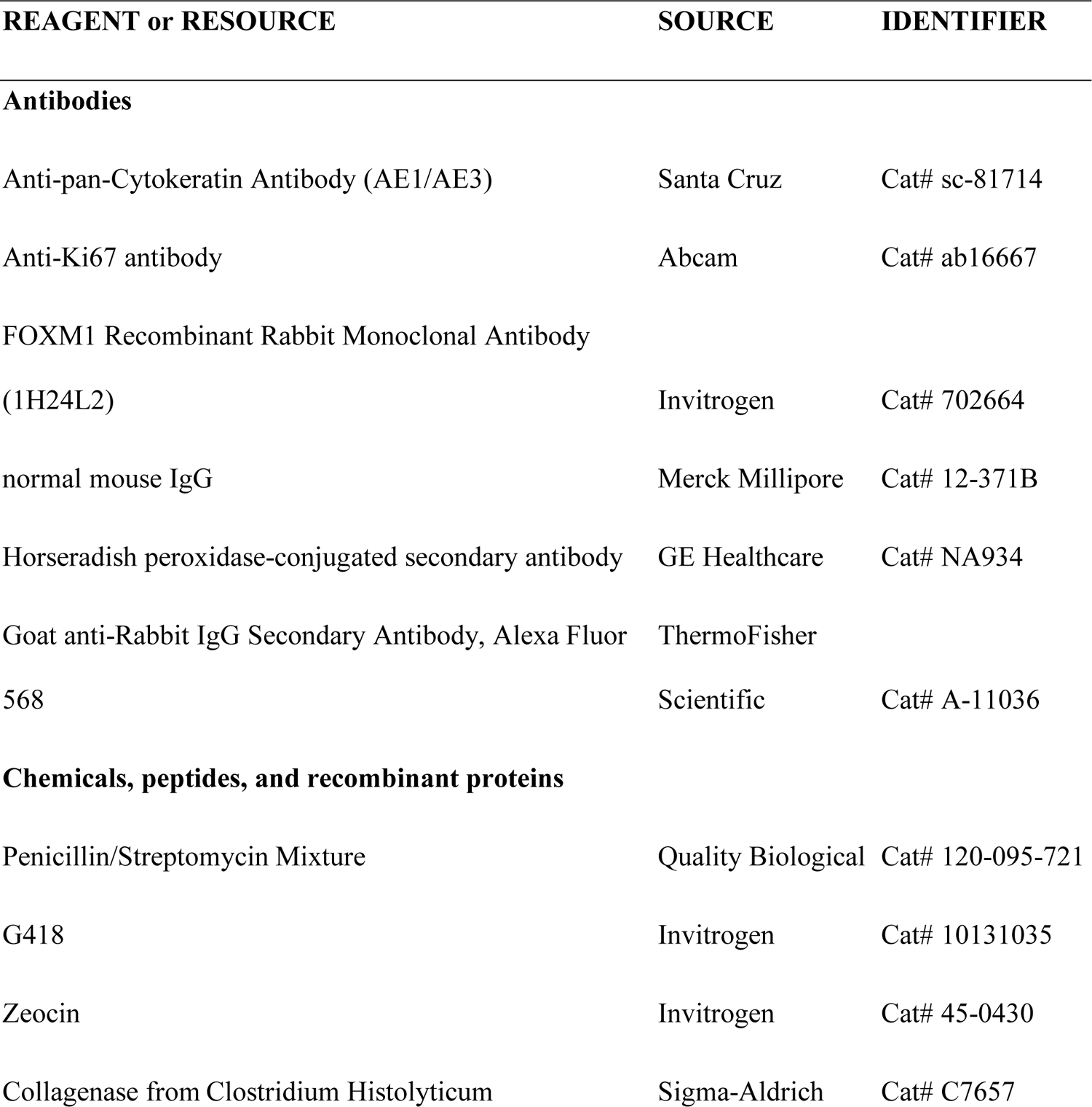
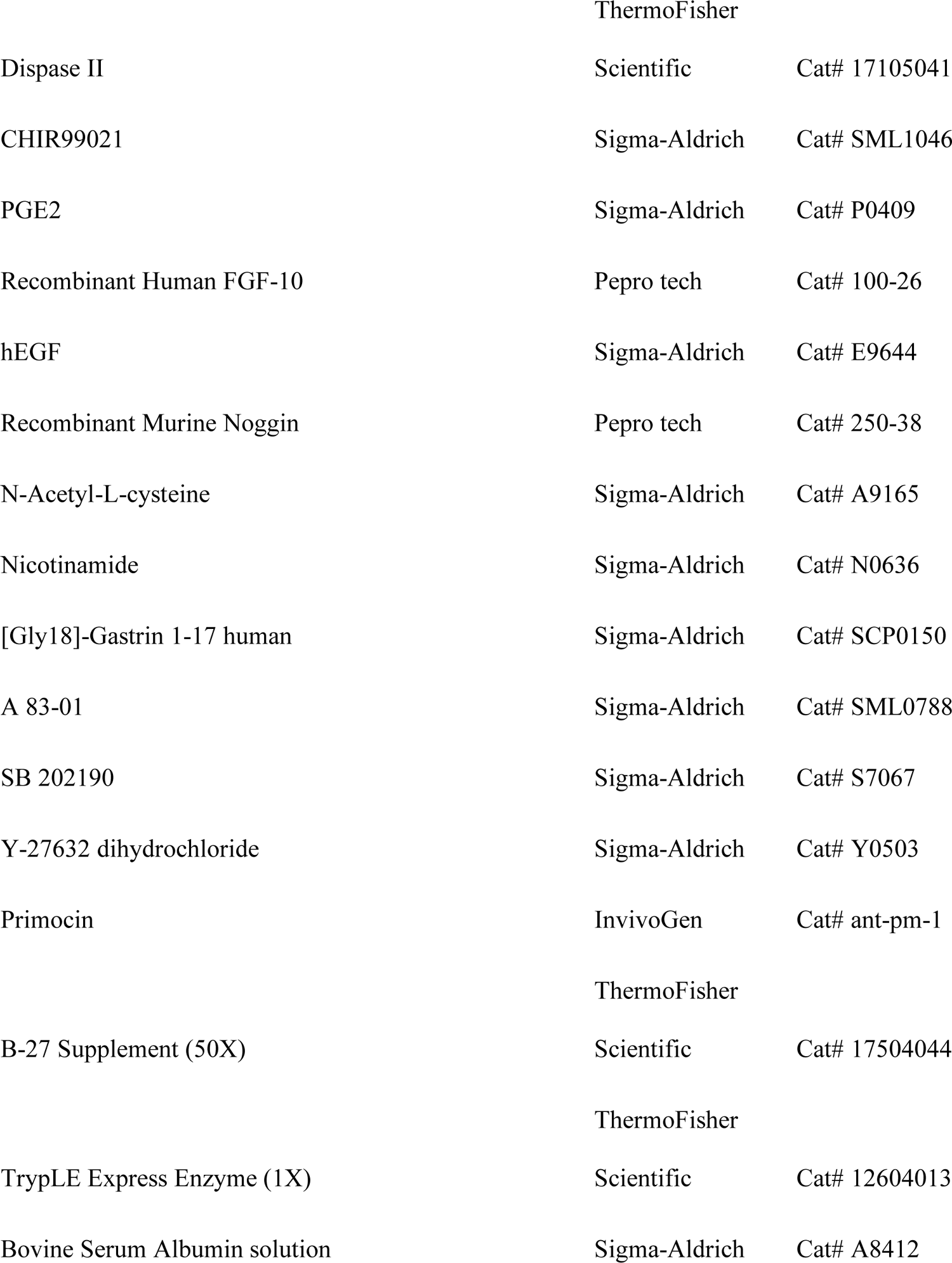
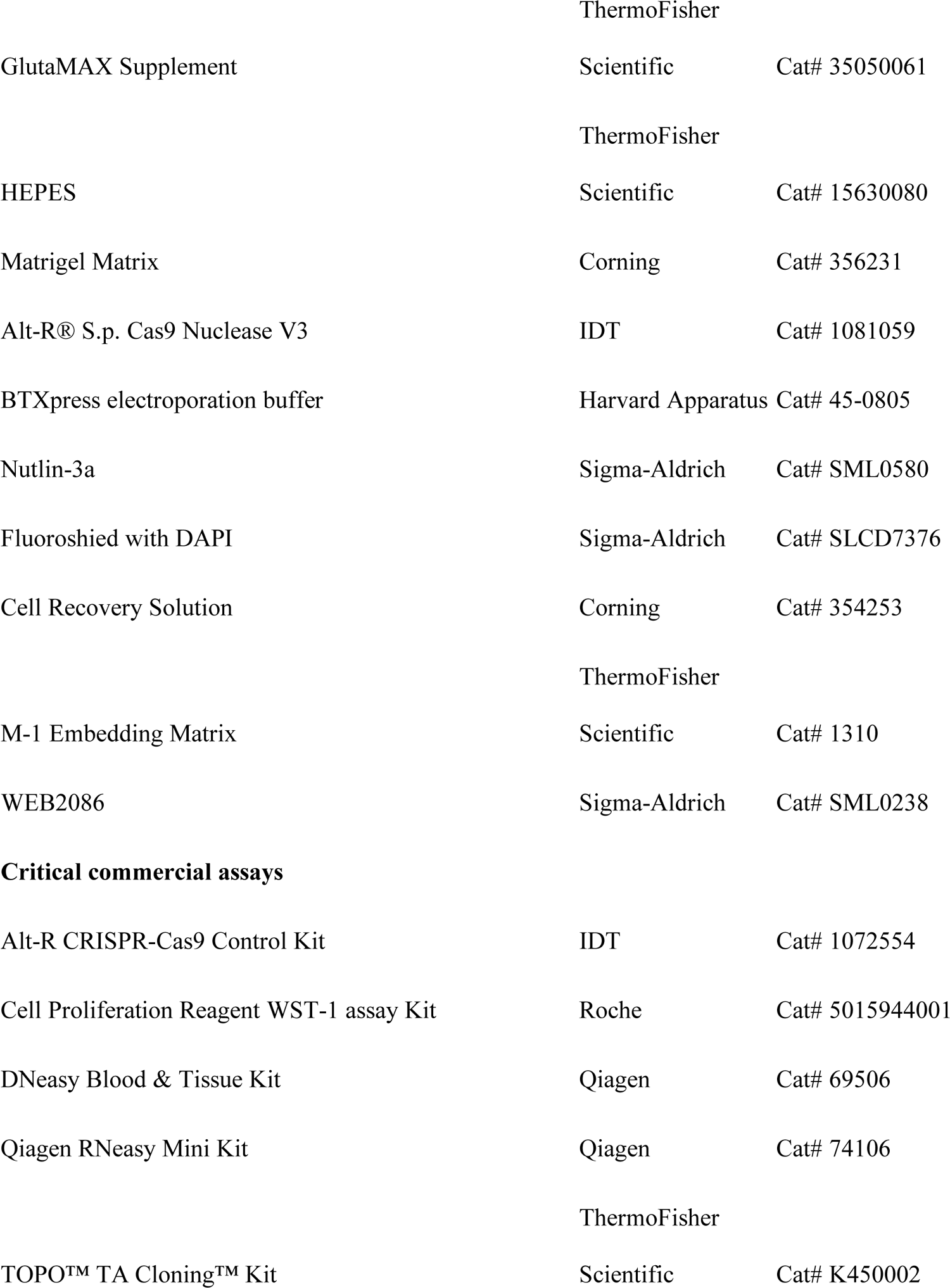
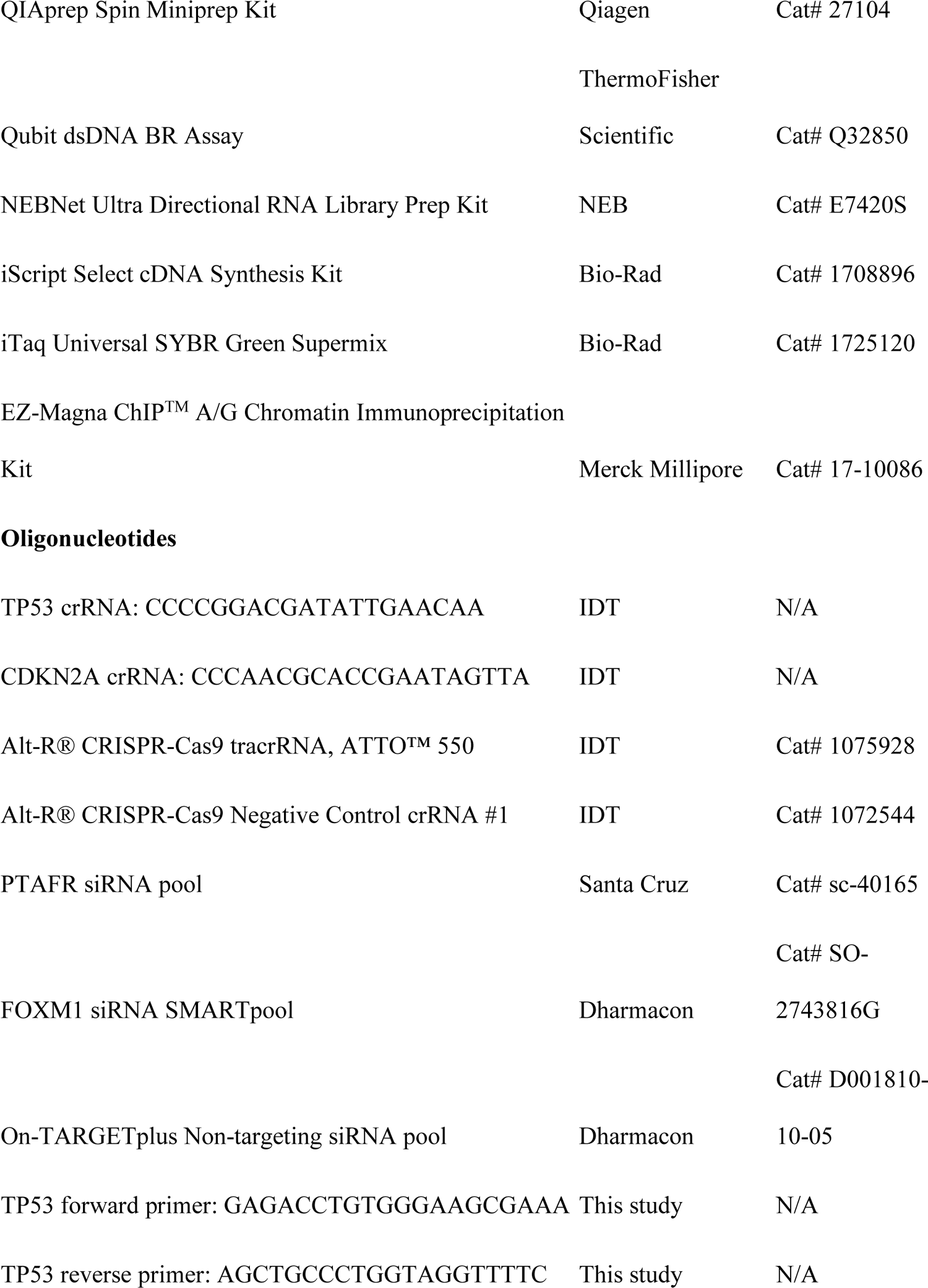
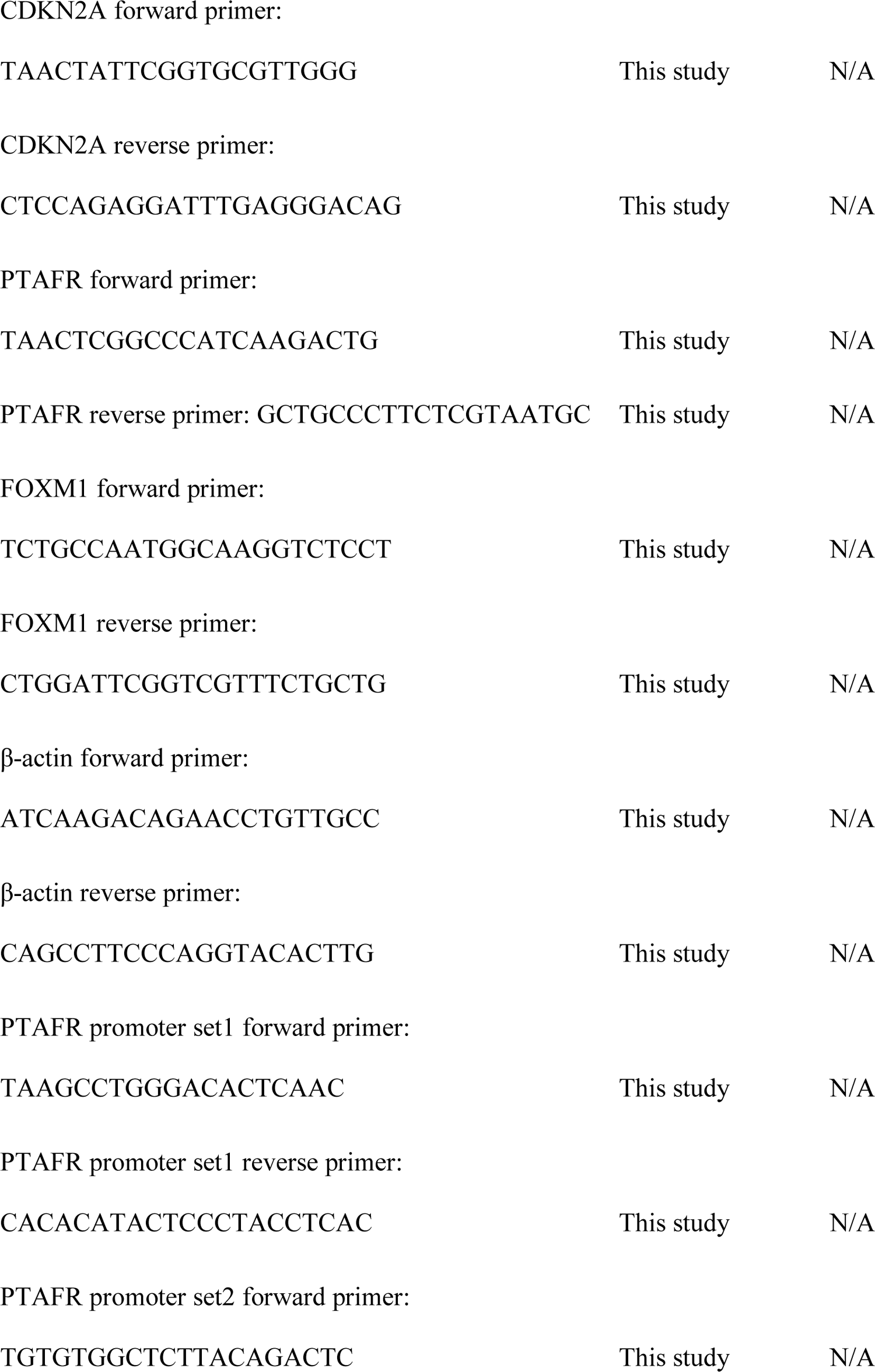
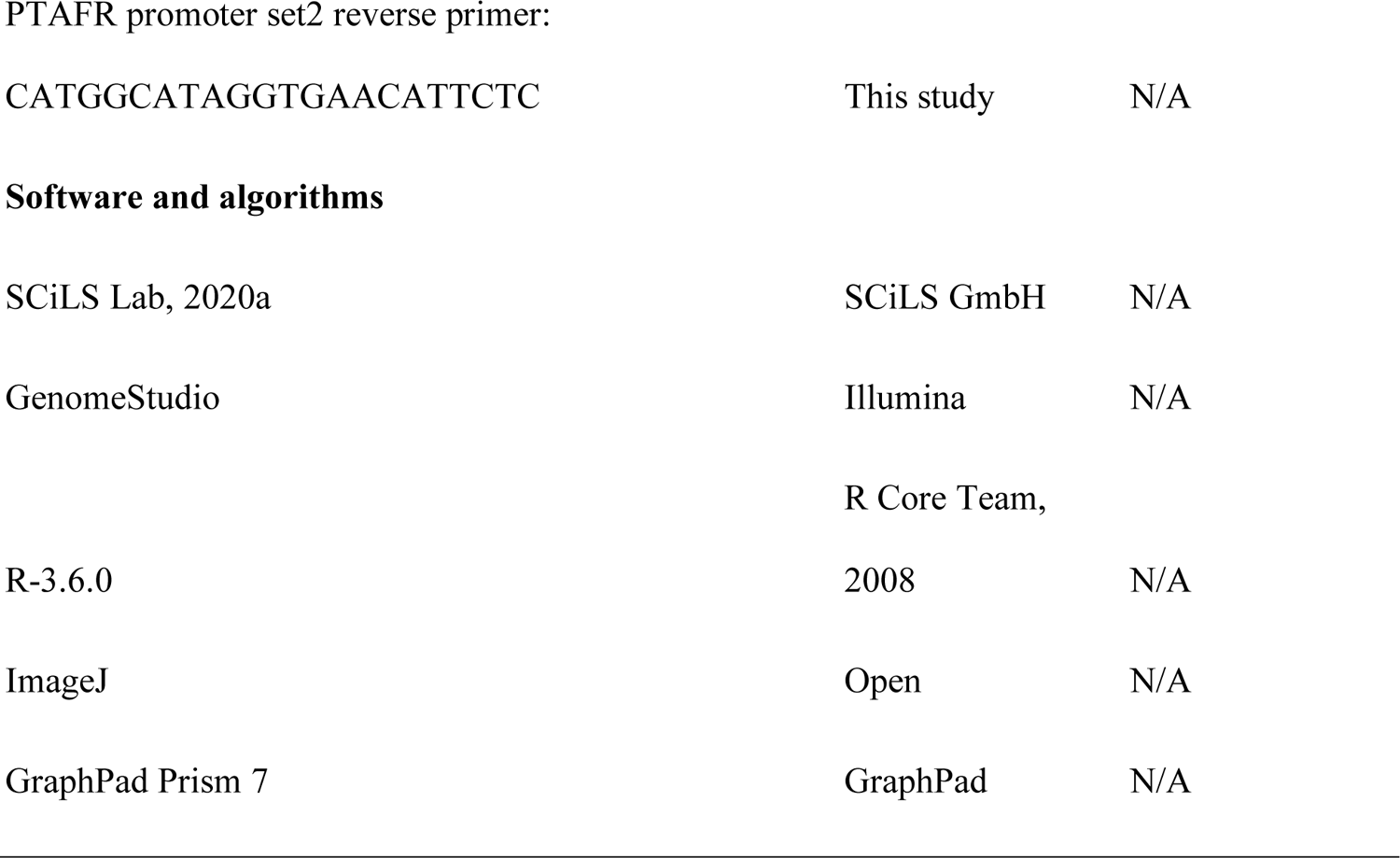
Key resources table.

### *TP53/CDKN2A* loss promotes proliferation, dysplasia, and neoplastic transformation of GEJ organoids

Next, we sought to generate a disease and pathologic model that could grow more vigorously, survive longer in culture, and better facilitate studies of GEJ neoplastic transformation. Because *TP53* and *CDKN2A* are two of the most frequently inactivated tumor suppressor genes in human GEJ cancers(*12–14*), we chose to inactivate these two genes in our GEJ organoid model using the CRISPR-Cas9 genome editing system. An all-in-one Cas9:gRNA ribonucleoprotein (RNP) complex targeting *TP53* (exon 4) and *CDKN2A* (exon 1α) was prepared using a gRNA complex (which combines crRNA and tracrRNA) and Cas9 nuclease. We then dissociated human GEJ organoids into small cell clusters (5 - 15 cells each) and delivered the RNP complex by electroporation using an optimized protocol (**Fig. S1A**). For the control organoid group, we electroporated a negative control, non-targeting RNP complex. A high electroporation efficiency of 42% was achieved (**Fig. S1B**). Subsequently, we selected for organoids with mutant-*TP53* using Nutlin-3a, which inhibits MDM2 and hence induces growth arrest of *TP53*-wildtype cells (*15*) (**Fig. S1C**). Upon selection, we performed Sanger sequencing to validate specific editing of targeted *TP53* and *CDKN2A* exons. Frameshift mutations, including 1-bp insertions or deletions at *TP53* and *CDKN2A* target sites (**Fig. 2A and B**), were observed, verifying successful genome-editing of GEJ organoids.

**Fig. 2.**
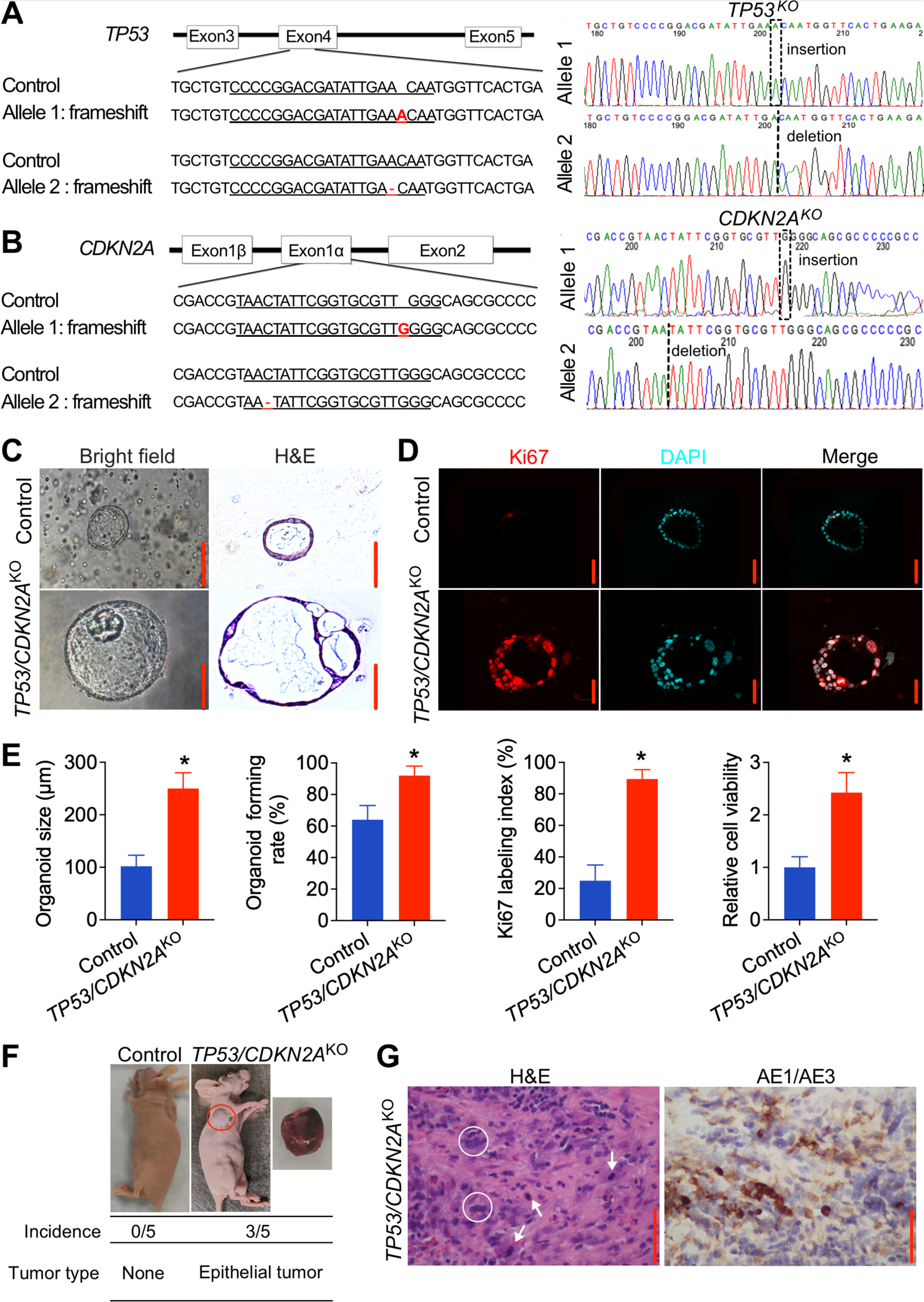
Knockout of *TP53/CDKN2A* promotes neoplastic transformation in human normal GEJ organoids. (A and B) Sanger sequencing of *TP53/CDKN2A*^KO^ GEJ organoids showing 1-bp insertion or deletion. *Red font* indicates corresponding frameshift indels in the genomic DNA. (C and D) On the 10th day after seeding 1x 10^5^ dissociated organoid cells, organoid cultures were photomicrographed using phase-contrast microscopy and collected for (C) H&E and (D) IF staining for Ki67 (red color). (E) Average organoid size, organoid-forming efficiency, and Ki67 index were determined by measuring > 50 organoids. *, P < 0.05. (F) Representative images of xenografts from mice injected with control or *TP53/CDKN2A*^KO^ GEJ organoids; the underlying table shows incidence and tumor characteristics. (G) Representative H&E and AE1/AE3 pan-keratin IHC staining (brown) in xenografts arising from *TP53/CDKN2A*^KO^ organoids. White arrows: mitoses; white circles: abnormally large, pleomorphic cells with irregular nuclear envelopes; scale bar = 100 µm.

We next characterized phenotypic changes of GEJ organoids upon loss of *TP53* and *CDKN2A*. Control organoids formed a single layer of epithelial cells with normal nuclei at day 10 after seeding. In sharp contrast, *TP53/CDKN2A*^KO^ organoids exhibited substantially larger diameters, more complex multicellular structures, increased mitotic activity, and markedly enlarged, atypical nuclei. These changes were consistent with dysplastic morphology (*16*) (**Fig. 2C and D**).

Organoid-forming rate also increased significantly in *TP53/CDKN2A*^KO^ relative to control organoids (92% *vs*. 64%, **Fig. 2E**). Immunofluorescence (IF) staining showed a striking elevation of Ki67 labeling index in *TP53/CDKN2A*^KO^ *vs.* control GEJ organoids (89.4% *vs.* 24.9%, **Fig. 2D and E**). Consistently, WST-1 assays revealed a 2.4-fold increase in proliferation of *TP53/CDKN2A*^KO^ *vs.* control organoids (**Fig. 2E**). Furthermore, under the same culture conditions, while control organoids could only be propagated for up to 6 months, *TP53/CDKN2A*^KO^ organoids were propagated for more than 19 months. These data demonstrate that loss of *TP53/CDKN2A* potently enhances proliferation and dysplasia of GEJ organoids *in vitro*.

To assess the *in vivo* effect of *TP53/CDKN2A* inactivation in GEJ organoids, we performed xenotransplantation assays. Control and *TP53/CDKN2A*^KO^ organoid cells (2×10^6^ cells/injection) were subcutaneously injected into the left and right armpit of 5 nude mice, respectively. Within a 5-month post-injection observation interval, no tumors were formed in 5 mice injected with control GEJ organoids. In contrast, *TP53/CDKN2A*^KO^ organoids developed tumors in 3 of 5 injected mice within 8 weeks (**Fig. 2F**). Importantly, these tumors appeared morphologically similar to highly differentiated gastroesophageal adenocarcinoma. H&E and IHC analysis of xenografts arising from *TP53/CDKN2A*^KO^ organoids showed increased mitosis, abnormally large, pleomorphic cells with irregular nuclear envelopes, and positive expression of AE1/AE3 proteins (**Fig. 2G**). Thus, these data *in vivo* suggest that inactivation of *TP53/CDKN2A* directly promotes neoplastic transformation in GEJ organoids.

### Lipidomic MALDI Imaging MS (IMS) identifies PTAFs as top upregulated lipids in *TP53/CDKN2A*^KO^ organoids

Reprogramming and dysregulation of lipid metabolism is a hallmark of cancer (*17*). However, it is unknown if and how lipid metabolic processes are altered during early GEJ carcinogenesis. To address this knowledge gap, we applied lipidomic MALDI-IMS to discover altered lipid species in *TP53/CDKN2A*^KO^ *vs.* control organoids using 2,5-dihydrobenzoic acid (DHB) as matrix in positive ion mode to ionize a broad range of phospholipids (*18*). Within the mass range from m/z 40 to 2,000, we obtained mass spectra of lipid species through direct analysis of organoid sections (**Fig. S2A**), while ion images were generated from each peak and displayed as the position in the organoid section and relative intensity (**Fig. S2B**). We applied a cutoff of m/z > 450 to minimize the background matrix signal in imaging experiments, as well as focus on phospholipids which are generally larger than m/z 400. We first performed analysis based on fold-changes of mean intensity data for individual peaks identifying 50 upregulated peaks and 132 downregulated peaks in *TP53/CDKN2A*^KO^ *vs.* control organoids under average fold-change > 1.5 (**Fig. 3A** and **Table S1**). In parallel, we performed receiver operating characteristic (ROC) analysis, using an area-under-the-curve (AUC) threshold value of > 0.75. By this criterion, 16 and 50 peaks were respectively upregulated and downregulated in *TP53/CDKN2A*^KO^ *vs.* control organoids (**Fig. 3B** and **Table S2**). Concordantly, all of these altered peaks identified by the ROC method were dysregulated in the same directions based on their fold-changes (**Fig. 3B**). We next performed on-organoid MS/MS (Fig. S3) and identified that among the 16 shared upregulated peaks, the one with the highest AUC (*m/z* 467.20) was a specific platelet-activating factor (PTAF) lipid PC-O-14:0 (<u>LMGP01020009</u>).

**Fig. 3.**
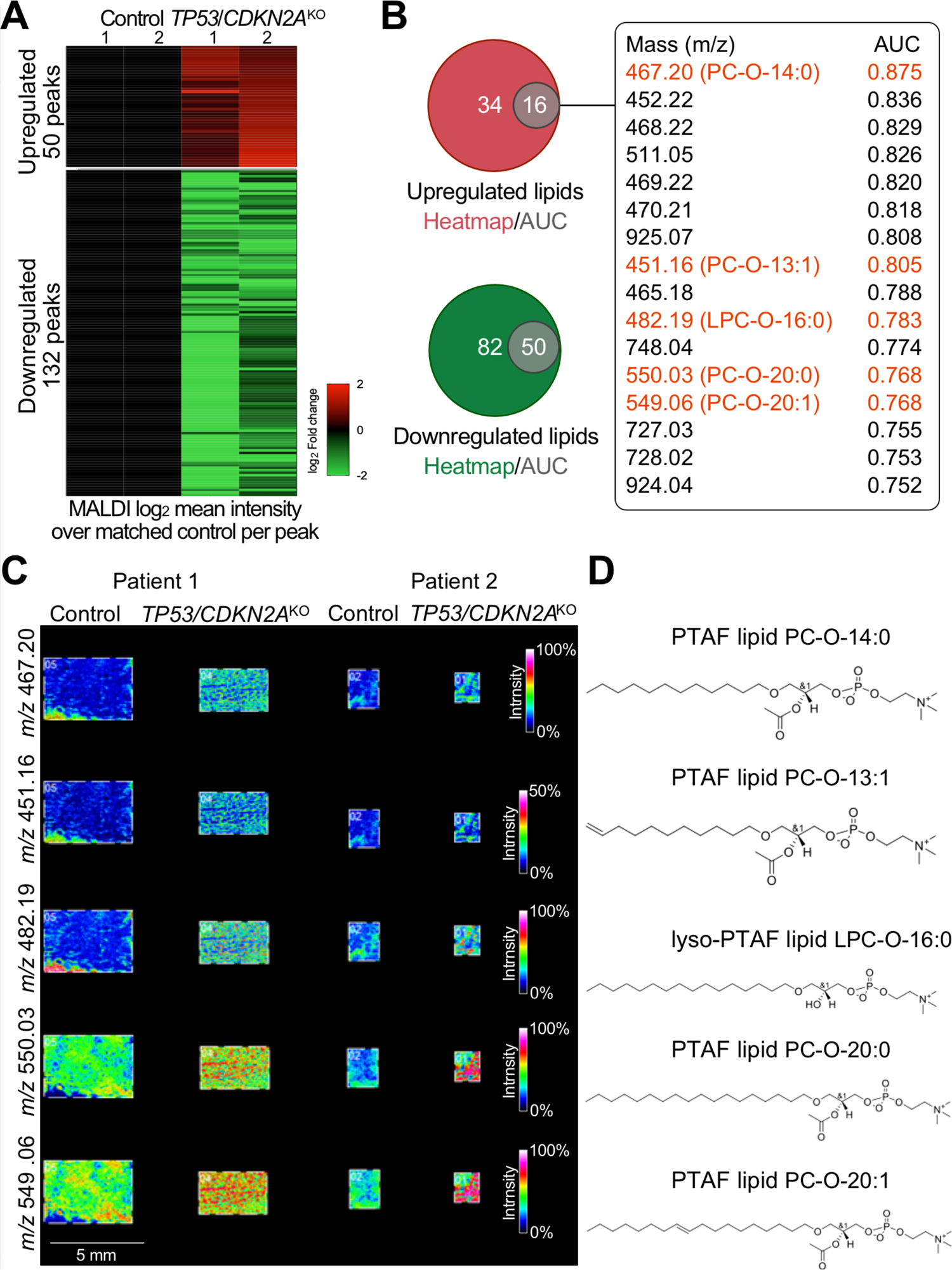
Platelet-activating factor (PTAF) lipids are notably increased phospholipids in *TP53/CDKN2A*^KO^ vs. control GEJ organoids. Matrix-assisted laser desorption/ionization (MALDI) imaging-based lipidomic analysis was performed on independent paired sets of *TP53/CDKN2A*^KO^ *vs*. control GEJ organoids derived from two different patients. (A) Heat map of discriminative lipid peaks (m/z) with a cutoff of m/z > 450, mean absolute fold change value > 1.5, and individual absolute fold change > 1.2 in each *TP53/CDKN2A*^KO^ organoids, compared with matched Control. (B) Venn diagrams represent the overlap of upregulated and downregulated lipids based on AUC and heatmap results. Overlapping 16 upregulated lipids and AUC values are listed. (C) MALDI imaging and (D) corresponding chemical structures of representative PTAF lipids in *TP53/CDKN2A*^KO^ organoids.

Moreover, 4 additional lipids were identified as either PTAF lipids (*m/z* 451.16, PC-O-13:1, <u>LMGP01020146</u>; *m/z* 550.03, PC-O-20:0, <u>LMGP01020094</u>; and *m/z* 549.06, PC-O-20:1, <u>LMGP01020146</u>) or a precursor of a PTAF lipid (*m/z* 482.19, LPC-O-16:0, <u>LMGP01060010</u>). MALDI imaging data and chemical structures of these lipids are displayed in **Fig. 3C** and **D**. Interestingly, PTAF lipids are a family of glycerophosphocholines implicated as bioactive mediators in diverse pathologic processes, including tumor angiogenesis and metastasis (*19, 20*).

### Inhibition of PTAF/PTAFR suppresses neoplastic properties of *TP53/CDKN2A*^KO^ GEJ organoids

Following identification of multiple PTAF lipids as the notably increased phospholipids in *TP53/CDKN2A*^KO^ GEJ organoids, we addressed their potential roles in GEJ neoplasia development. As a glycerophosphocholines, PTAFs exert biological effects by binding to their cognate receptor, PTAFR (*21*). Thus, we first evaluated PTAFR levels in GEJ neoplasia, both in our organoid model and in The Cancer Genome Atlas (TCGA) dataset. Interestingly, similar to its cognate lipid ligand, PTAFR expression was upregulated in *TP53/CDKN2A*^KO^ *vs.* control organoids (**Fig. 4A**). Consistent with this finding, TCGA EAC tumors exhibited higher PTAFR mRNA levels than did nonmalignant TCGA GEJ samples (**Fig. S4**). Encouraged by these observations, we next directly abrogated PTAF/PTAFR function in our early GEJ model system by either siRNA knockdown or pharmacologic inhibition. Importantly, silencing PTAFR expression by siRNAs significantly decreased the average size, cell viability, and Ki67 index of *TP53/CDKN2A*^KO^ (**Fig. 4B-E**). In parallel, we treated *TP53/CDKN2A*^KO^ organoids with either vehicle control (0.1% DMSO) or a specific PTAFR pharmacologic antagonist, WEB2086, at various concentrations. In agreement with the siRNA results, *TP53/CDKN2A*^KO^ organoids displayed significantly reduced size and Ki67 index after WEB2086 treatment (**Fig. 4F to H**). WST-1 assays showed that metabolically active cells began to decrease on day 4, and a time- and dose-dependent inhibitory effect was confirmed on day 7 and 10 (**Fig. 4H**).

**Fig. 4.**
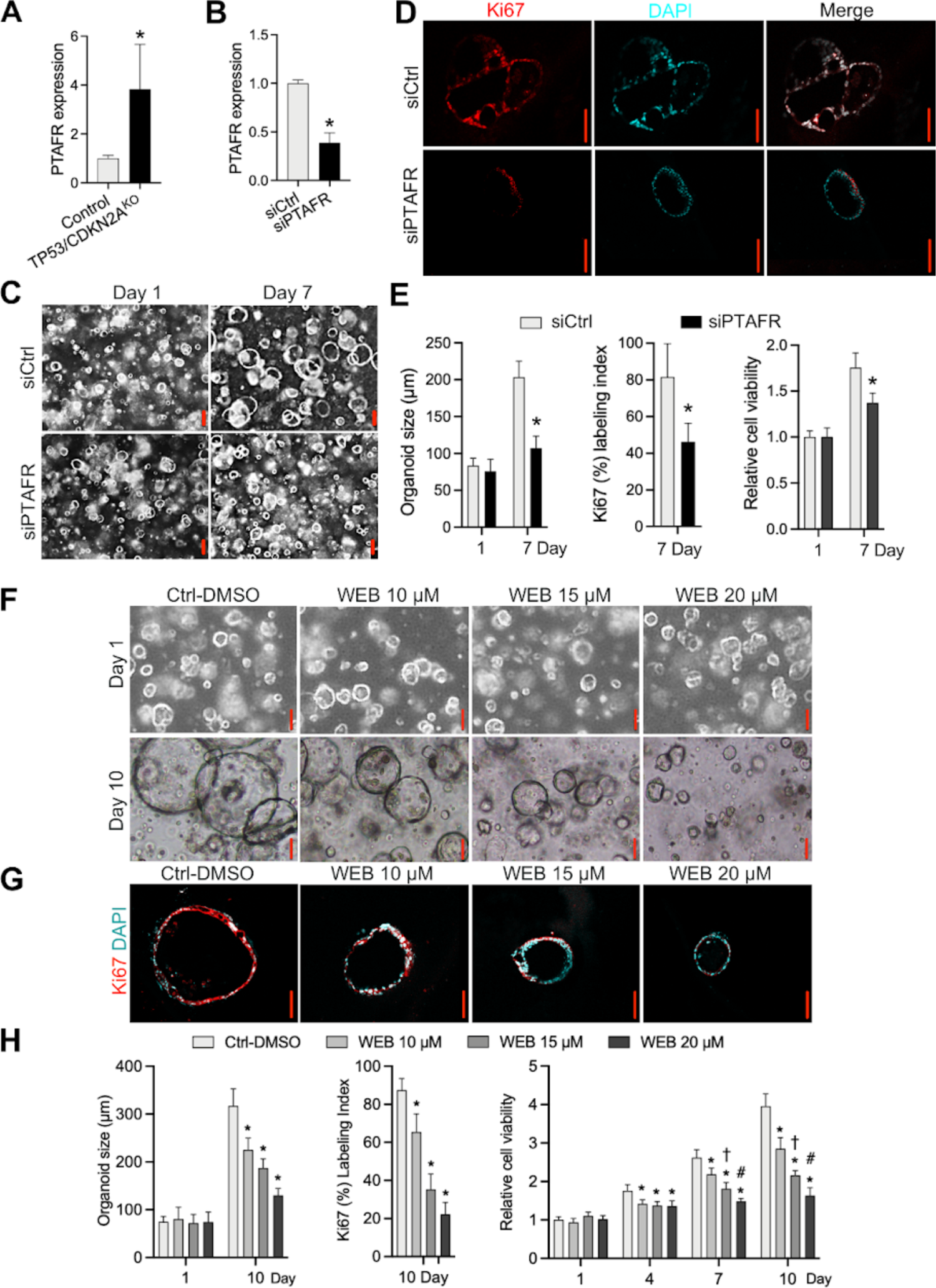
Blockade of PTAF/PTAFR inhibits growth and proliferation of *TP53/CDKN2A*^KO^ GEJ organoids. (A) mRNA levels of PTAFR in *TP53/CDKN2A*^KO^ *vs.* control GEJ organoids. (B) Knockdown of PTAFR mRNA by siRNAs in *TP53/CDKN2A*^KO^ GEJ organoids. (C) Average organoid size and (E) cell viability determined by phase-contrast imaging and WST-1 assays, respectively. (D and E) Ki67 IF images and quantification obtained on Day 7. *, P < 0.05 *vs.* same-day siCtrl. Scale bar = 100 µm. (F-H) **A**verage organoid size and cell viability (H) determined by phase-contrast imaging and WST-1 assays, respectively. (G and H) Ki67 labeling images and quantification obtained on Day 10. Scale bar = 100µm. *, P < 0.05 *vs.* Ctrl-DMSO on the same day; †, P < 0.05 *vs.* WEB 10 µM on the same day; #, *P* < 0.05 *vs.* WEB 15 µM on the same day.

Next, we tested the inhibition of PTAFR *in vivo. TP53/CDKN2A*^KO^ organoid cells (2×10^6^ cells/injection) were subcutaneously injected into the armpits of nude mice. Mice were treated with vehicle control (1.25% DMSO) or WEB2086. Within a 3-month post-injection observation interval, 3 out of 5 injected mice developed tumors in the DMSO control group within 7 weeks (**Fig. 5A**), similar to what we observed above (**Fig. 2F**). Importantly, WEB2086 treatment completely prevented tumor formation in *TP53/CDKN2A*^KO^ organoids (**Fig. 5A**). We next assessed the effect of WEB2086 in a xenograft model derived from an established EAC cell line (Eso26). Again, PTAFR inhibition resulted in significant suppression of EAC xenograft growth and proliferation (**Fig. 5B and C**). IHC analysis showed downregulated expression of Ki67 in WEB2086-treated Eso26 tumor xenografts (**Fig. 5D**). These results demonstrate that blockade of the PTAF/PTAFR lipid cascade potently inhibits *TP53/CDKN2A*^KO^ GEJ organoid growth, proliferation, and tumorigenesis *in vitro* and *in vivo*, suggesting an important function for PTAF/PTAFR in mediating early neoplastic progression at the GEJ.

**Fig. 5.**
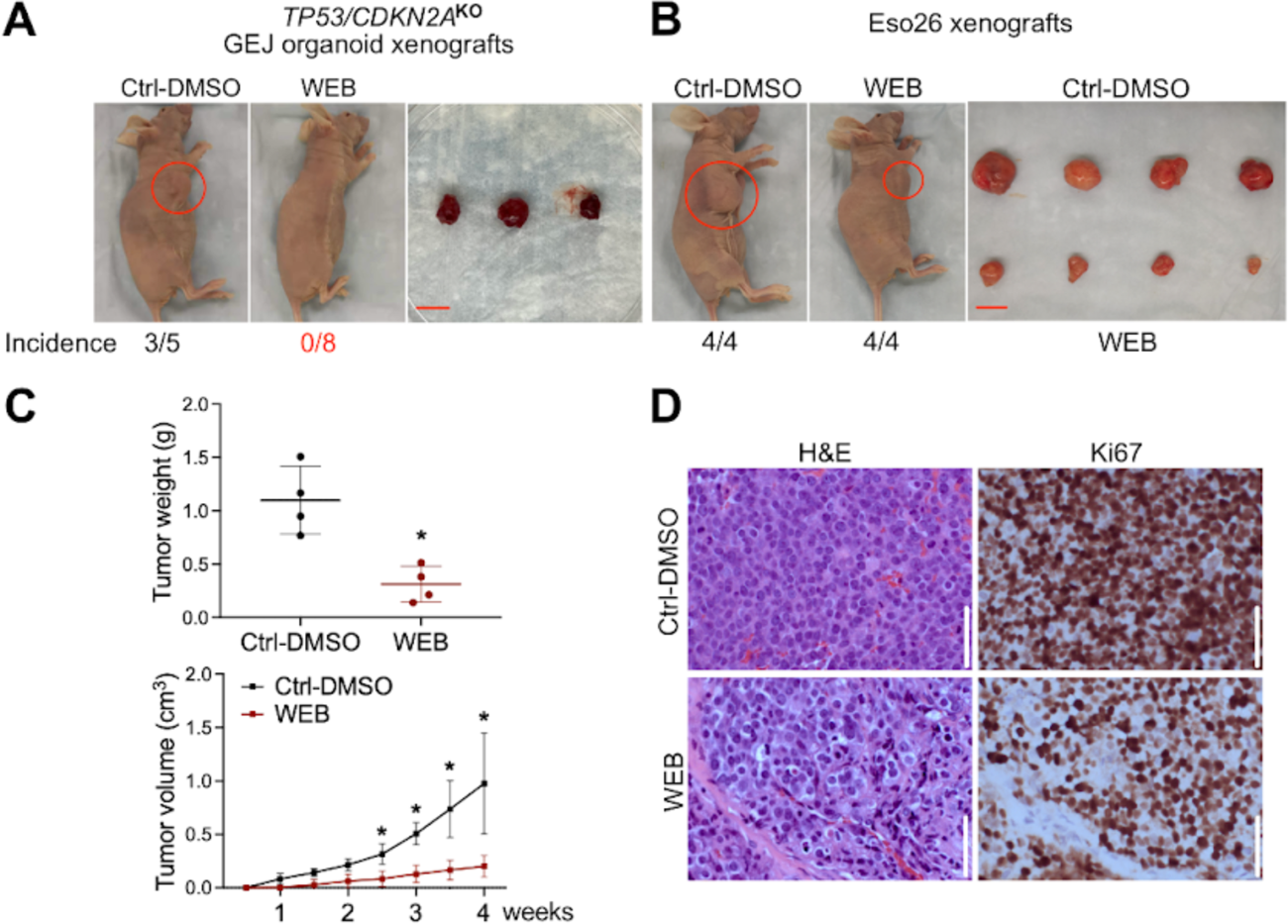
Blocking the PTAF/PTAFR pathway inhibits *in vivo* tumorigenesis of *TP53/CDKN2A*^KO^ GEJ organoids and Eso26 cells. (A) Xenograft images and incidence rates after WEB2086 treatment in *TP53/CDKN2A*^KO^ GEJ organoids. Scale bar = 1 cm. (B) Xenograft images and incidence rates after WEB2086 treatment in Eso26 cells. Scale bar = 1 cm. (C) Significant inhibition of tumor growth and weight of Eso26 xenografts by WEB2086. *, P < 0.05. (D) Ki67 IHC and hematoxylin-eosin staining in Eso26 xenografts. Scale bar = 100 µm.

### Inactivation of *TP53/CDKN2A* alters the DNA methylome and transcriptome of GEJ organoids

To explore differences at the transcriptomic and epigenomic levels between control and *TP53/CDKN2A*^KO^ organoids, we performed transcriptome sequencing (RNA-seq) and Illumina Methylation EPIC array profiling, respectively. Specifically, paired wild-type and *TP53/CDKN2A*^KO^ organoids from 4 patients were subjected to RNA-seq. Compared with the control group, *TP53/CDKN2A*^KO^ organoids contained 556 significantly differentially expressed genes (312 upregulated and 244 downregulated; **Fig. 6A** and **Table S3**). Gene Ontology analysis of these genes identified strong enrichment of biological processes and pathways related to mitotic entry and cell cycle progression (**Fig. 6B** and **Table S4**). This finding is consistent with the above observation of significantly accelerated proliferation and growth of *TP53/CDKN2A*^KO^ organoids. At the epigenomic level, we determined both differentially methylated CpGs and differentially methylated regions (DMRs) between control and *TP53/CDKN2A*^KO^ organoids derived from each of the 4 patients. For example, in organoids derived from patient 1, a total of 1,732 and 1,391 CpG sites were significantly hypomethylated in the mutant and control organoids, respectively (**Fig. 6C**). These CpG sites corresponded to 129 and 83 hypomethylated DMRs in *TP53/CDKN2A*^KO^ and control organoids, respectively (**Fig. 6D** and **Table S5**). Results of organoids derived from the other three patients are shown in **Fig. S5** and **Table S5**. DNA hypomethylated regions are known to contain regulatory elements associated with the binding of transcription factors (TFs) (*22, 23*). To identify candidate TFs implicated in our neoplastic GEJ organoid model, we investigated enriched TF-recognition motif sequences in hypomethylated DMRs using the HOMER package (*24*). Notably, motifs of the Forkhead box (FOX) TF family were among the most enriched sequences in hypomethylated DMRs in *TP53/CDKN2A*^KO^ organoids (**Fig. 6E**). Because different FOX TF family members recognize similar motif sequences (*25*), we next sought to identify which factor(s) were involved. Previous studies have shown that expression levels of a particular TF correlate with levels of demethylation in regions it occupies (*26, 27*). Therefore, we analyzed expression levels of the 26 FOX TFs based on RNA-seq data from our organoids, as well as on normal GEJ and EAC data from the TCGA. Both FOXM1 and FOXC2 were strongly increased in *TP53/CDKN2A*^KO^ organoids. FOXM1 was also upregulated in EAC *vs*. normal GEJ tissues (**Fig. 6F**), suggesting it as a candidate TF enriched in hypomethylated DMRs with possibly increased activity in *TP53/CDKN2A*^KO^ organoids. FOXM1 is a known regulator of cell proliferation and cell cycle progression in cancer (*28, 29*), in line with the pro-neoplastic phenotypes in our *TP53/CDKN2A*^KO^ organoids.

**Fig. 6.**
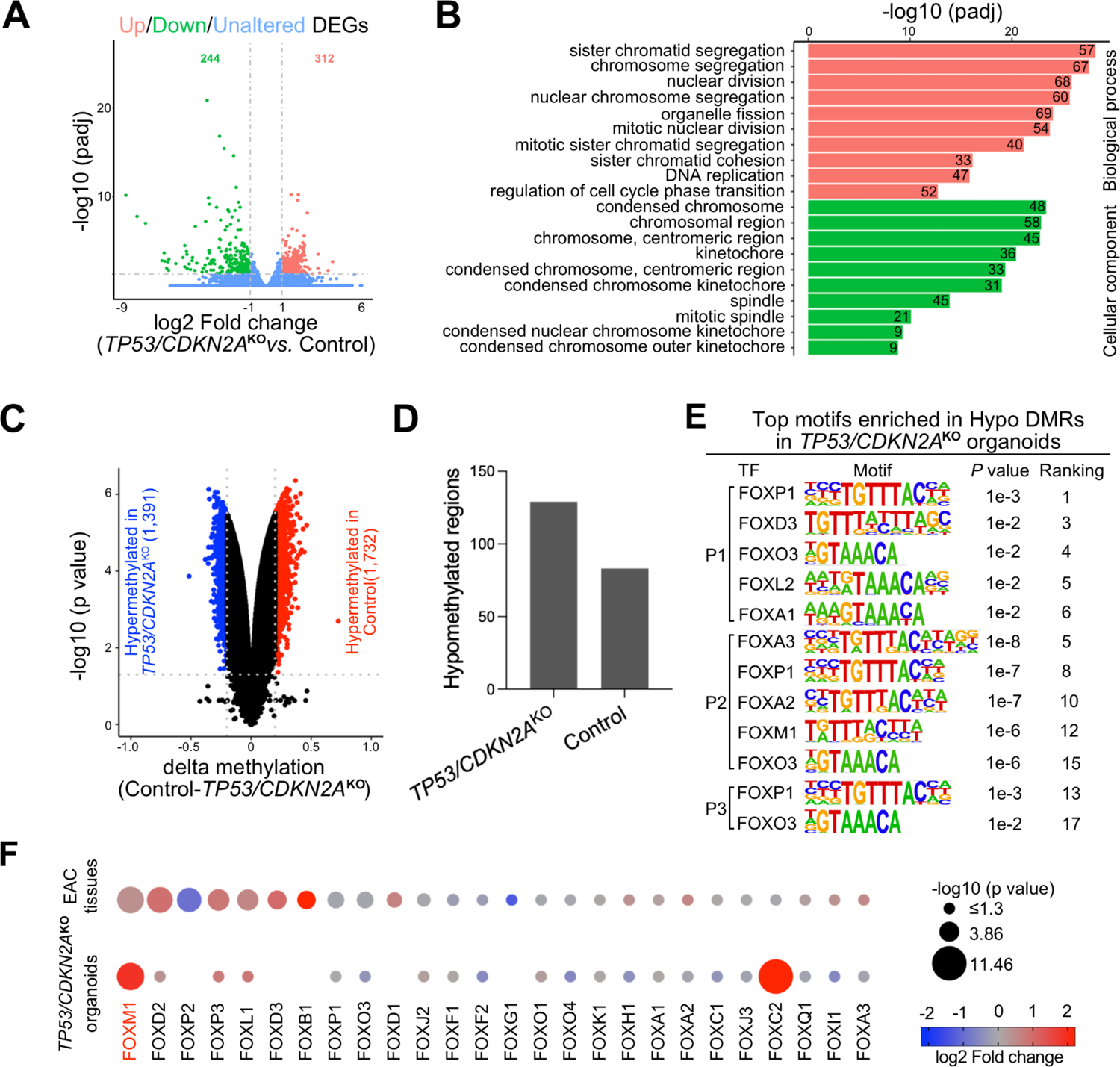
DNA methylome and transcriptome profiling of *TP53/CDKN2A*^KO^ GEJ organoids. (A) A volcano plot showing differentially expressed genes (DEGs) in *TP53/CDKN2A*^KO^ organoids *vs*. control organoids. (B) Gene ontology (GO) analysis identified top-ranked key terms of biological processes and cellular components enriched in DEGs in *TP53/CDKN2A*^KO^ *vs*. control organoids. (C) A volcano plot showing differentially methylated CpGs and (D) column plots showing DMRs in T*P53/CDKN2A*^KO^ vs. control organoids. (E) Top-ranked TF binding motifs from the Forkhead box (FOX) family enriched in hypomethylated DMRs in *TP53/CDKN2A*^KO^ *vs.* control organoids. *P* values were calculated using the HOMER package. (F) Dot plots showing expression levels of FOX family TFs in *TP53/CDKN2A*^KO^ *vs.* control organoids (bottom dots) and in EAC tumors *vs.* normal GEJ samples from the TCGA (upper dots). Genes are ranked based on the -log10 (p-value) comparing EAC tumors with normal GEJ samples. The dot size denotes - log10 (p-value); color indicates log2 fold change; missing dots correspond to undetectable mRNA expressions.

### PTAFR is a direct transcriptional target of FOXM1

Our above data not only identified PTAFs phospholipids as one of the most induced classes of lipid molecules in our lipidomic profiling, but also revealed the upregulation of its cognate receptor, PTAFR, in *TP53/CDKN2A*^KO^ GEJ organoids (**Fig. 4**). To further elucidate mechanisms mediating upregulation of PTAFR expression in *TP53/CDKN2A*^KO^ organoids, we performed motif enrichment analysis of the PTAFR promoter, which revealed that the FOX motif sequence again was highly enriched (No.4) (**Fig. 7A**). This was particularly interesting given our above data identifying FOXM1 as a potential regulator involved in epigenetic alterations in *TP53/CDKN2A*^KO^ organoids.

**Fig. 7.**
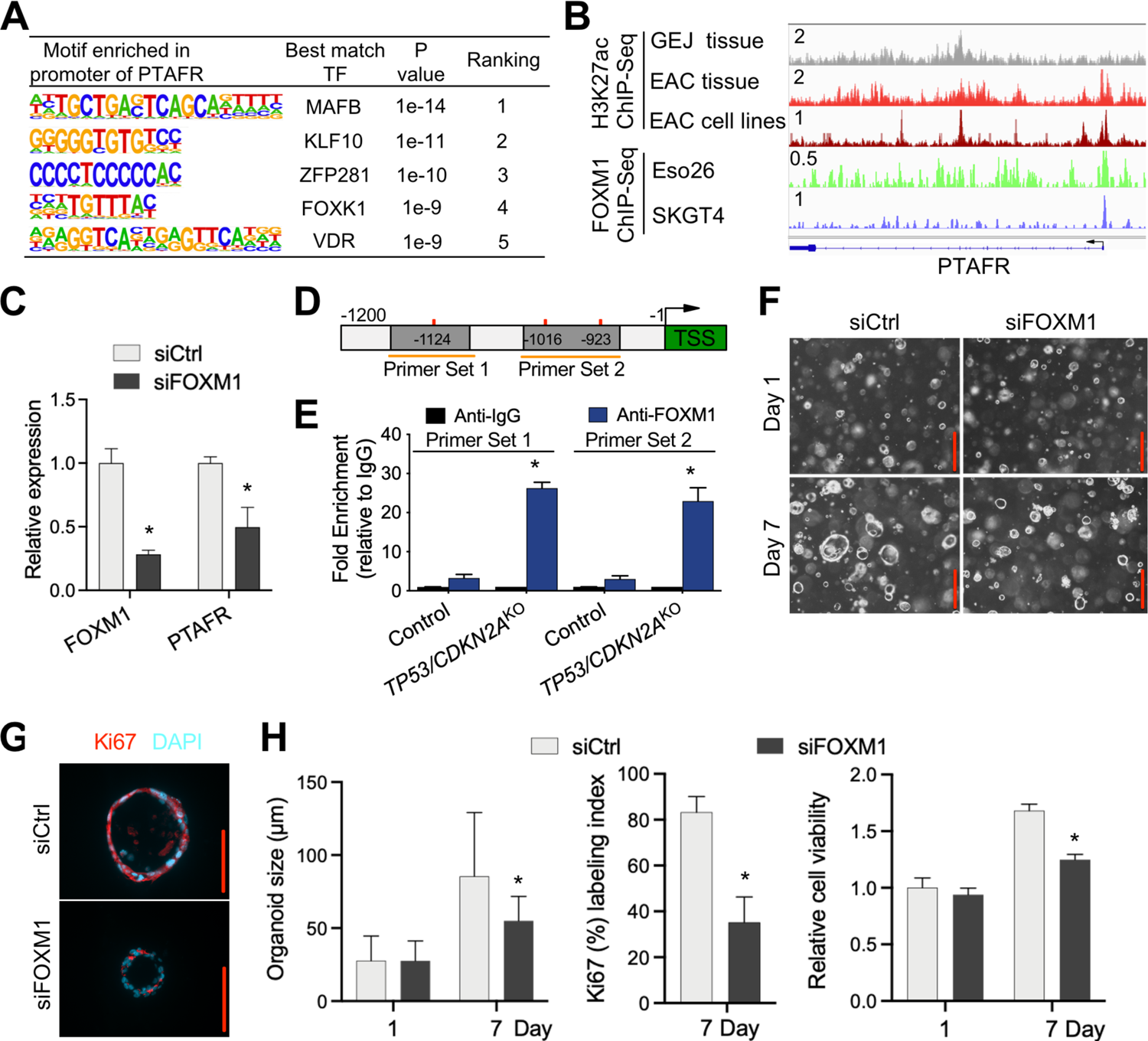
PTAFR is a direct downstream target of FOXM1. (A) Motif enrichment analysis of PTAFR promoter region. (B) ChIP-seq profiles for H3K27ac and FOXM1 at the PTAFR locus indicated tissues and cell lines. All H3K27ac signals are shown at the same scale. (C) Knockdown of FOXM1 by siRNAs reduced PTAFR mRNA transcript levels in *TP53/CDKN2A*^KO^ GEJ organoids. (D) Schematic of PTAFR transcriptional start site (TSS) (*green*) and upstream sequence. PCR primers for ChIP experiments (*orange horizontal lines*) and FOXM1 binding motifs (*red vertical lines*) are indicated. (E) ChIP-qPCR detection of FOXM1 occupancy at the *PTAFR* promoter region, using primer sets indicated in (D). ChIP was performed with either anti-FOXM1 or anti-IgG antibodies, and fold enrichment relative to anti-IgG is shown. (F-H) Average organoid size determined by phase-contrast imaging and cell viability determined by WST-1 assays, respectively. Ki67 labeling images and quantification obtained on Day 7. Scale bar = 100µm. *, P < 0.05 *vs*. siCtrl on the same day.

To validate whether FOXM1 regulates the transcription of PTAFR, we first performed ChIP-seq using an anti-FOXM1 antibody. Indeed, FOXM1 occupied the promoter of *PTAFR* in two different EAC cell lines (**Fig. 7B**). Moreover, our recent H3k27ac ChIP-seq data (*30*) showed that *PTAFR* harbors strong H3k27ac signals at its promoter and candidate enhancers in both EAC primary tumors and cell lines (**Fig. 7B**), suggesting robust transcriptional activation. In contrast, in normal GEJ samples, H3k27ac was barely deposited at the *PTAFR* locus, indicating weak/inactive transcription (**Fig. 7B**). This pattern of histone modification was in agreement with expression levels of PTAFR, which was upregulated in EAC *vs.* normal GEJ samples (**Fig. S4**).

Since the above ChIP-seq data were generated in EAC cell lines and primary tumors, we next performed FOXM1 ChIP-qPCR in GEJ organoids. Importantly, endogenous FOXM1 showed prominent occupancy at the *PTAFR* promoter in *TP53/CDKN2A*^KO^, with a more than 7.9-fold increase in enrichment compared to wild-type GEJ organoids (**Fig. 7 D and E**). To validate the regulation of FOXM1 upon PTAFR transcription, we silenced FOXM1 via siRNA in *TP53/CDKN2A*^KO^ organoids, which showed that FOXM1 knockdown significantly reduced PTAFR expression (**Fig. 7C**). In addition, FOXM1 silencing reduced average organoid size, cell viability, and Ki67 index in *TP53/CDKN2A*^KO^ organoids (**Fig. 7F to H**), phenocopying the effects we obtained with PTAF/PTAFR blockade. These data demonstrate that FOXM1 binds directly to the PTAFR promoter, thereby augmenting PTAFR expression and proliferation in *TP53/CDKN2A*^KO^ organoids.

## DISCUSSION

A major hurdle in understanding the molecular origins and biology of GEJ cancer is a paucity of appropriate biologically relevant models, particularly of early neoplastic events. Primary three-dimensional organoid cultures derived from normal epithelia represent a desirable system for studying critical properties of the original native tissue *in vitro*, including morphological, histological, and molecular features (*3*). To our knowledge, we describe here the first human normal GEJ-derived organoid culture model, with a highly reliable protocol ensuring successful organoid culture from endoscopic biopsies. We show that wild-type GEJ organoids can be propagated *in vitro* for at least 4 months, and *TP53/CDKN2A*^KO^ organoids even longer (for at least 19 months). This platform offers great promise for modeling GEJ-associated diseases, characterizing healthy and diseased GEJ conditions, and discovering novel molecular mechanisms underlying the transition from normal to pathological GEJ.

Furthermore, we also demonstrate here that CRISPR-engineered human *TP53/CDKN2A*^KO^ organoids offer a productive tool for modeling early neoplastic events at the GEJ. *TP53* mutations have been detected in benign Barrett’s esophagus (BE) patients who later develop high-grade dysplasia or EAC (*31–33*) suggesting their involvement in early neoplasia at the GEJ anatomic region. Likewise, *CDKN2A* is inactivated in more than 80% of BE and EAC tissues (*34–36*). Similar molecular abnormalities also occur in gastric cardia neoplasia (*37–42*), supporting the contribution of *TP53* and *CDKN2A* to diverse types of adenocarcinogenesis involving the GEJ region. In agreement with these static observations, our dynamic model establishes that *TP53/CDKN2A* inactivation directly causes biologic and molecular features consistent with GEJ neoplastic progression.

*TP53* and *CDKN2A* have been implicated in several lipid metabolic processes (*43–46*). However, to our knowledge, direct evidence does not yet exist linking *TP53* and *CDKN2A* to lipid metabolic aberrations occurring during human GEJ carcinogenesis. Here, we demonstrate that *TP53/CDKN2A* inactivation directly alters lipidomic profiles in GEJ organoids. We discover that the lipids most upregulated by *TP53/CDKN2A*^KO^ in GEJ organoids include several PTAFs, a family of phospholipid mediators with known signaling effects in important biological processes, including cancer initiation and (*21, 47–50*). PTAF phospholipid levels are elevated in a feline model of esophagitis (*51*); Elevated PTAF phospholipid levels are significantly more frequent in gastric cancer than in control patients (*52*). PTAFs function primarily by binding to its cognate receptor, PTAFR (*53–55*). Notably, PTAFR is abundantly expressed in human gastric adenocarcinoma tissues; moreover, PTAFR overexpression is strongly associated with poor prognostic clinical parameters (*56–58*). Consistent with these known clinical and biological data, we identify here PTAF/PTAFR as an etiologic mediator in neoplastic progression induced by *TP53/CDKN2A*^KO^ at the GEJ.

Our study also uncovers the therapeutic potential of pharmacological PTAF/PTAFR inhibition during GEJ neoplastic progression. Although to date no PTAF/PTAFR-related cancer treatments are available, *in vitro* and *in vivo* experimental models have shown inhibition of tumorigenesis and cancer angiogenesis by PTAFR antagonists, including ginkgolide B, BN-50730, and WEB2170 (*20, 59*). In our model, the PTAFR antagonist WEB2086 inhibited early neoplastic changes in *TP53/CDKN2A*^KO^ organoids, while causing analogous growth suppression in ESO26 esophageal adenocarcinoma cells. Further research is warranted to assess the therapeutic prospect of PTAF/PTAFR inhibition in GEJ tumors.

Our data also suggest that *TP53*/*CDKN2A*^KO^ elicits extensive epigenetic and transcriptional reprogramming that propel the normal GEJ toward a malignant state. Here, our integrative analyses identify prominent enrichment of FOXM1 binding sequences in demethylated regions coupled with the upregulation of this TF, directly caused by *TP53*/*CDKN2A* inactivation. FOXM1 is overexpressed in multiple solid tumors (*60–62*), and signaling downstream of this TF contributes to cancer development and progression via cross-talk with multiple cell signaling pathways. Notably, we also demonstrate a novel mechanistic link between FOXM1 and PTAFR: FOXM1 binds to the PTAFR gene promoter, directly activating the transcription of PTAFR. Consistent with results upon PTAF/PTAFR inhibition, FOXM1 knockdown in *TP53*/*CDKN2A*^KO^ GEJ organoids leads to severe cell growth inhibition.

In summary, the human primary benign GEJ organoid model and its pro-neoplastic induction by *TP53/CDKN2A* knockout now enables molecular deconstruction of early GEJ tumorigenesis and neoplastic progression; furthermore, our lipidomic, epigenetic, and transcriptional profiling studies yield valuable insights into mechanistic underpinnings of GEJ malignancy. The highly induced phospholipid family, PTAFs, and their receptor, PTAFR, together show strong pro-neoplastic activity in GEJ evolution while simultaneously suggesting potential therapeutic strategies against GEJ cancers. Conceivably, our strategies for transforming organoids derived from human primary normal epithelia into more neoplastic entities are generalizable to modeling other cancer-driving events, as well as other types of malignancy, where they may expand our understanding of tumor-specific gene-regulatory networks and yield novel potential therapeutic targets.

## MATERIALS AND METHODS

### Patient samples

In accordance with approved Institutional Review Board protocols at the Johns Hopkins Hospital, primary human endoscopic biopsy samples were acquired at the Johns Hopkins Hospital under written informed consent. Tissue samples were pathologically confirmed as nondysplastic GEJ.

### Cell lines and maintenance

L Wnt-3A cells (CRL-2647) were purchased from ATCC and maintained in DMEM-10% FBS to produce Wnt-3A-conditioned medium. Cultrex HA-R-Spondin1-Fc 293T cells (3710-001-01) were purchased from Bio-techne and maintained in DMEM-10% FBS to generate R-spondin-1-conditioned medium. Eso26 cells were purchased from JENNIO Biological Technology and grown in RPMI 1640-10% FBS.

### GEJ organoid cultures

GEJ organoids were established as described in Fig. 1A. Briefly, fresh endoscopic GEJ biopsies were preserved in ice-cold conditioned PBS (PBS containing 10 µM ROCK inhibitor Y27632, 2% (vol/vol) penicillin/streptomycin, and 1x Primocin) until further processing within 24 hours. After washing biopsies with conditioned PBS at least 5 times, samples were minced into fragments < 1 mm^3^ using microdissecting scissors. Tissue fragments were digested in DMEM containing 2.5% (vol/vol) FBS, 1% (vol/vol) penicillin/streptomycin, 1 mg/ml collagenase type IX, and 120 µg/ml dispase type II at 37 °C with 200 rpm shaking for 40-90 mins. Following centrifugation at 400 g at 4°C for 3 minutes, pelleted cell clusters were resuspended in Matrigel. Using 24-well plates, 2,000 cells were seeded per well in 50 µl of Matrigel. After incubating at 37°C for 10 minutes to solidify the Matrigel, 500 µl of growth medium was added to each well. The growth medium for GEJ organoids was Advanced DMEM/F12 supplemented with 50% (vol/vol) Wnt-3A conditioned medium (home-made), 20% (vol/vol) R-spondin-1 conditioned medium (home-made), 1% (vol/vol) penicillin/streptomycin, 10 nM PGE2, 100 ng/ml human rFGF-10, 50 ng/ml hEGF, 100 ng/ml Noggin, 1 mM N-acetylcysteine, 10 mM Nicotinamide, 10 nM Gastrin I, 500 nM A-83-01, 10 µM SB202190, 10 µM Y27632, 5 µM CHIR99021 (only for the first 1-2 passages), 1X Primocin, and 1X B-27 supplement. Culture medium was changed once every three days until the organoids were ready to passage. For passaging, organoids were washed in PBS and digested with TrypLE containing 10 µM Y27632 for 5–7 mins at 37°C. After incubation, DMEM/F12 was added to stop digestion. Organoids were mechanically dissociated by pipetting and centrifuged at 500 g for 3 mins. After resuspending the pellet in Matrigel, 50-100 µl per droplet of the cell-Matrigel suspension were plated onto a new culture plate. Reagents used for organoid culture are listed in the Table 1.

### Organoid viability assay (WST-1 assay)

To quantify metabolically active viable cells, organoids were seeded onto 96-well plates and cultured. At indicated time points, 10 µl per well of Cell Proliferation Reagent WST-1 assay kits were added to the 96-well plates and incubated with organoids for 90 minutes. After incubation, only media was transferred to the wells of a new 96-well plate, which was read at an absorbance of 450 nm by a Thermo Scientific Microplate Reader. All experiments were performed in triplicates.

### CRISPR-Cas9 genomic editing of GEJ organoids

Organoids were electroporated using the NEPA21 (Nepa Gene) system and the Alt-R CRISPR-Cas9 System (IDT). Cas9:gRNA ribonucleoprotein (RNP) complex was prepared as follows: to make the 100 µM gRNA complex, 200 µM tracrRNA labeled with ATTO™ 550 and 200 µM crRNA were mixed in equimolar concentrations, heated at 95°C for 5 mins, and then allowed to slowly cool to room temperature; to produce the RNP complex for each electroporation, 6 µl of gRNA complex (100 µM), 8.5 µg of Cas9 Nuclease (10 µg/µl), and 10.5 µl of Duplex Buffer were combined and incubated at room temperature for 10 mins. RNP complex was stored for further use at −80°C. Two days before electroporation, organoids were passaged and maintained in organoid culture medium w/o antibiotics including 5 µM CHIR99021. Organoids were dissociated into clusters of 10-15 cells, resuspended in 80 µl of Electroporation Buffer containing 4 µM Electroporation Enhancer, and then mixed with 25 µl of RNP complex targeting *TP53* and 25 µl of RNP complex targeting *CDKN2A*. The mixture was transferred into a precooled 2-mm electroporation cuvette. Electroporation parameters were set according to Fujii et al (*63*). After electroporation, 400 µl of prewarmed culture medium including 5 µM CHIR99021 was immediately added to the electroporation cuvette. Cells were seeded after incubation for 40 mins at 37℃. Two days after electroporation, transfection efficiency was measured by fluorescence microscopy. Three days after electroporation, organoids were treated with 10 µM Nutlin-3a for functional selection of *TP53-*mutant cells for 2-3 weeks, on the basis that Nutlin-3a inhibits the proliferation of *TP53* wild-type cells. Organoids electroporated with the negative control RNP complex were used as the control group. To validate targeted mutations, genomic DNA from edited organoids was extracted, followed by PCR amplification, TOPO-cloning and Sanger sequencing. All reagents and sgRNA sequences used in this section are provided in the Table 1.

### MALDI mass spectrometric imaging of the lipidome in GEJ organoids

Organoids were transferred to even molds and immersed in M-1 Embedding Matrix after being isolated from Matrigel using Cell Recovery Solution and washed with cold PBS for 3 times. Organoid molds were wrapped in aluminum foil and floated on liquid nitrogen for progressive freezing. Frozen organoids were equilibrated to −20 °C, cryosectioned at 10-µm thickness and thaw-mounted onto temperature-equilibrated, hexane-and-ethanol-washed indium tin oxide (ITO) slides (Delta Technologies, Loveland, CO) on a Leica CM1860 UV cryostat (Wetzlar, Germany). All organoids were sectioned in a layout that maximizes the number of sections per slide to compare *TP53/CDKN2A*^KO^ *vs.* control GEJ organoids. Several serial sections were cryosectioned with the same layout for technical repeats. Slides were sprayed with 40 mg/mL DHB dissolved in 70% HPLC-grade methanol/30% HPLC-grade water using an HTX-M5 sprayer (Chapel Hill, NC) with the following parameters: nozzle temperature −75 °C, 8 passes, 0.1 ml/min flow rate, 1200 mm/min nozzle velocity, 3 mm track spacing, criss cross (CC) pattern, 10 psi pressure, 3 l/min gas flow rate, and 10 second drying time. The final matrix density was 8.89 × 10^-3^ mg/mm^2^ and the linear flow rate was 8.33 × 10^-5^ ml/mm.

Matrix-assisted laser desorption/ionization (MALDI) mass spectrometry imaging was acquired on Bruker MALDI TOF/TOF rapifleX instrument (Bruker Daltonik, Bremen, Germany) in the Johns Hopkins Applied Imaging Mass Spectrometry (AIMS) Core in reflectron-positive mode, at 20-micron pixel size with a 20-micron raster and 20-micron imaging laser, 200 laser shots per pixel, and a mass range of m/z 40 to 2,000. Imaging data was collected using FlexImaging version 5.0 and FlexControl version 4.2 (Bruker Daltonik, Bremen, Germany) and data was normalized to total ion current (TIC) for all data analysis). MALDI lipid imaging data were imported into SCiLS Lab software (Version2020a, SCiLS GmbH, Bremen, Germany) and alignment was performed with TIC to remove any drift artifacts from imaging. Quantitative spectral, pixel-based, paired comparisons (*64–66*) between *TP53/CDKN2A*^KO^ and control GEJ organoids. For structural identification, m/z’s of the top lipids were identified by on-tissue MS/MS using collision-induced decay (CID) with argon using the single beam laser with a resultant field of 54 x 54 microns with 4000 laser shots and an isolation window of ± 2 Da. MS/MS spectra were collected from both KO organoids. For initial identification, the Lipid Maps Structure Database (LMSD) (*67*) was used by uploading a peak list, searching [M+H]^+^ and [M+Na]^+^ with a mass tolerance of ± 0.2 m/z and all lipid classes selected. This generated a list of potential hits which were used to solve MS/MS spectra in ChemDraw Professional version 16.0 (PerkinElmer, Waltham, MA).

### Genome-wide DNA methylation profiling and data analysis

DNA methylation profiles for 4 paired sets of control and double-knockout GEJ organoids were generated using the Illumina Methylation/EPIC array platform, which combines bisulfite conversion of genomic DNA and whole-genome amplification with direct, array-based capture and scoring of CpG loci. Genomic DNA was extracted from organoids using DNeasy Blood & Tissue kits. All DNA samples were quantified by Qubit dsDNA BR Assay, assessed for purity by A260/280 and A260/230 ratio, and examined for integrity by electrophoresis on 0.8% agarose gels. DNA samples were then hybridized to Infinium Methylation EPIC BeadChips, following the Infinium HD Methylation Assay Protocol (*68*).

The SeSAME package (*69*) was used to extract the DNA methylation value of each probe using the openSesame function. Recommended general masking probes were removed according to the annotation file of Infinium DNA methylation arrays. Differentially methylated probes were identified by the limma package (version 3.46.0) with adjusted p-value < 0.05, absolute delta methylation change > 0.2. We further identified differentially methylated regions (DMRs) based on differentially methylated probes by the DMRcate package, with Fisher’s exact test p-value < 0.05.

### RNA-sequencing (RNA-Seq) and data analysis

RNA sequencing was performed on 4 paired sets of control and double-knockout GEJ organoids derived from 4 patients. Total RNA was extracted and treated with DNase I before sequencing. Libraries were constructed using NEBNet Ultra Directional RNA Library Prep kits. Quantified libraries were sequenced on the Illumina NovaSeq 6000 platform, and paired-end reads were generated. An index of the reference genome was built and paired-end clean reads were aligned to the reference genome using Hisat2 v2.0.5. Read counts for each gene were generated by FeatureCounts v1.5.0-p3. FPKM of each gene was calculated based on the length of the gene and the read count mapped to this gene. DEseq2 results were used for differential expression analysis, and genes with an adjusted p-value < 0.05 and an absolute fold change > 2 found by DESeq2 were designated as differentially expressed. Gene Ontology (GO) enrichment analysis of differentially expressed genes was implemented using the clusterProfiler R package (*70*).

### Real-Time Quantitative PCR

Total RNA was extracted using RNeasy kits, and DNA was eliminated via on-column DNase digestion. 500 µg RNA was reversely transcribed using iScript Select cDNA Synthesis kits (Bio-Rad). Quantitative PCR was performed using iTaq Universal SYBR Green Supermix. All results were normalized to β-actin expression. Primers used for qRT-PCR are listed in the Table 1.

### siRNA-directed gene silencing

Two days prior to electroporation, organoids were passaged and cultured in organoid culture medium without antibiotics including 5 µM CHIR99021. Organoids were dissociated into clusters of 10-15 cells, resuspended in 100 µl of Electroporation Buffer containing 4 µM Electroporation Enhancer, and then mixed with 10 µl of 50 mM siRNA. The organoids were electroporated using the same procedure mentioned above 24 hrs after electroporation. Total DNA from cloned organoids was then extracted and reversely transcribed into cDNA, followed by qRT-PCR to validate knockdown efficiency. siRNA sequences are provided in the Table 1.

### Histology, immunofluorescence and immunohistochemistry

Organoid cultures and tissues were fixed overnight in 10% formalin at room temperature. Paraffin-embedded organoids and tissues were serially sectioned into 10 µm slices. Paraffin sections were deparaffinized, rehydrated, followed by either staining for hematoxylin and eosin (H & E), or antigen retrieval in sub-boiling 10 mM sodium citrate buffer pH 6.0 for 10 mins. For immunofluorescence (IF), slides were permeabilized in 0.5% TritonX-100 in PBS and blocked in 1% goat serum in PBS for 30 mins at room temperature. After blocking, slides were incubated with anti-Ki67 (1:200, Abcam) overnight in a humidified chamber at 4 °C. Sections were washed by PBST (3 times for 5 mins each) and incubated with Alexa Fluor secondary antibodies (1:500) for 1 hr. After washing with PBST, slides were mounted with Fluoroshied with DAPI (Sigma). Immunohistochemistry was performed using the automated Bond-Max autostainer by Leica for the following antibodies: AE1/AE3 (1:200, Santa Cruz). Images were acquired with a Lecia Inverted Confocal SP8 (Johns Hopkins Medicine Ross Imaging Center). Reagents used here are listed in the Table 1.

### Chromatin immunoprecipitation (ChIP)

ChIP experiments were performed using EZ-Magna ChIP^TM^ A/G Chromatin Immunoprecipitation Kit (Merck Millipore, 17-10086) according to the manufacturer’s procedures. Cells were crosslinked with 1% formaldehyde and lysed with cell lysis buffer containing 1X Protease Inhibitor Cocktail II. Nuclei were isolated with nuclear lysis buffer supplemented with 1X Protease Inhibitor Cocktail II. The chromatin extract was sonicated (8 mins total, AmpL 30%, pulse on 10 s, pulse off 20 s) and sheared to a length between 200 bp to 1,000 bp on wet ice. The sheared crosslinked chromatin was immunoprecipitated with antibodies co-incubated with magnetic protein A/G beads. Antibodies included anti-FOXM1 (5 µg per CHIP reaction, Invitrogen, 702664) and normal mouse IgG (1 µg per reaction). Purified DNAs were subjected to qPCR. qPCR primers are listed in the Table 1.

### Xenotransplantation in nude mice

All procedures and experimental protocols involving mice were approved by the Animal Experimental Committee of the Johns Hopkins University School of Medicine. For xenotransplantation of organoids, 2×10^6^ organoid cells or ESO26 cells resuspended in cold 50% Matrigel were injected into the axillary of nude mice. For the WEB2086 treatment assay, 5 mg/kg.d of WEB2086 or vehicle control (1.25% DMSO in PBS) was administered by intraperitoneal injection every two days for three weeks. Xenograft size was measured twice a week. The xenograft volume (V) was monitored by measuring the length (L) and width (W) with a caliper based on the formula V=1/2×(L×W^2^). Mice were sacrificed at the end of the experiment, and xenografts were excised for further analyses.

## Statistical analysis

The data are presented as mean ± SD unless indicated otherwise. Statistical analysis was assessed using GraphPad Prism 9.2. For in vitro experiments, we used student’s t-test or one-way ANOVA unless otherwise noted in the figure legends. For xenograft experiments, we used unpaired Student’s t-test. P < 0.05 was considered statistically significant.

## List of Supplementary Materials

Fig. S1 to S5 for multiple supplementary figures

Table S1 to S5 for multiple supplementary tables

## Acknowledgments

We are indebted to the patients who participated in the study. We also thank Hans Clevers and Jarno Drost for key reagents. We thank Pro. Bon-Kyoung Koo, George McNamara, Dr. Qi Cao (University of Southern California), and Dr. Kevin Waters (Cedars-Sinai Medical Center) for expert technical assistance. We would like to acknowledge the Johns Hopkins Applied Imaging Mass Spectrometry (AIMS) Core Facility at the Johns Hopkins University School of Medicine.

## Author contributions

Conceptualization: HZ, YC, D-CL, SJM

Methodology: HZ, YC, AK, KM, Y-YZ, BZ, CT, KG, EJS, SN, MK, VS, RAA, SJ, NW, WC, XL

Investigation: HZ, YC, AK, KM, Y-YZ, BZ, CT, KG, EJS, SN, MK, VS, RAA, SJ, WC

Visualization: HZ, Y-YZ, BZ, CT, KG, RAA Funding acquisition: D-CL, SJM

Project administration: YC, D-CL, SJM Supervision: D-CL, SJM

Writing – original draft: HZ, AK Writing – review & editing: All authors

## Data and materials availability

Reagents, resources and materials employed in this study are detailed in the Table 1. Materials and reagents used in this study and generated at our laboratory are available upon request. RNA-seq datasets of TCGA EAC and normal GEJ tissues can be obtained from XENA data portal (http://xena.ucsc.edu/). All the other data supporting the findings of this study are available within the paper and its supplementary information files. Further information and requests for resources and reagents should be directed to and will be fulfilled by the Lead Contacts, De-Chen Lin and Stephen J. Meltzer.

---

**Figure S1.**
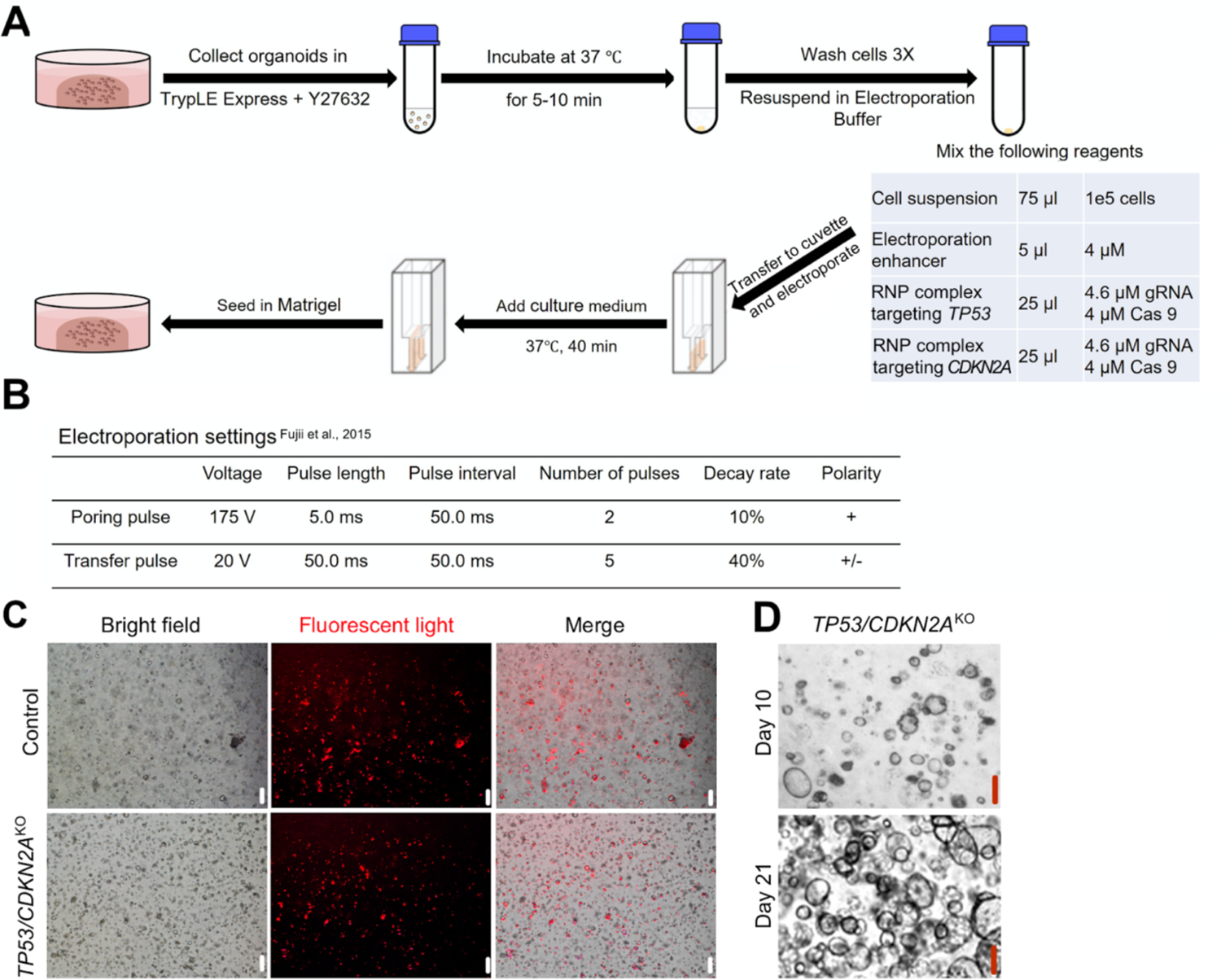
*TP53/CDKN2A*^KO^ GEJ organoids created by CRISPR-Cas9 genome editing. (A) Electroporation preparation workflow and (B) two-step electroporation settings. (C) Cas9 nuclease and the negative control or *TP53/CDKN2A*-targeted gRNA complex were successfully transfected into GEJ organoids (red fluorescence). (D) survival and sustained growth of *TP53/CDKN2A*^KO^ GEJ organoids in Nutlin-3a-containing selective medium. *Scale bar = 100 μm*.

**Figure S2.**
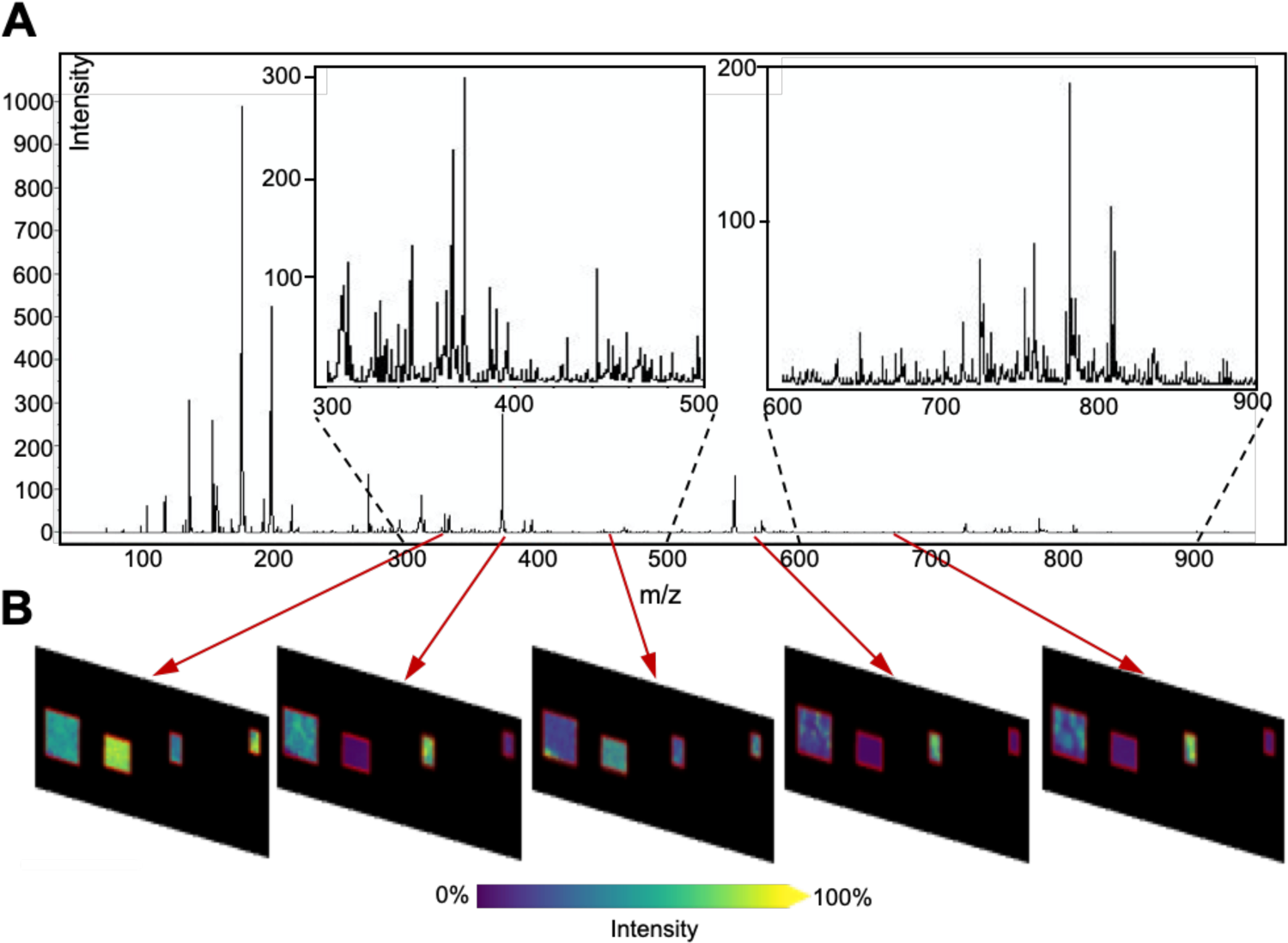
Average lipid spectra and MALDI images of control and *TP53/CDKN2A*^KO^ GEJ organoids. (A) Average lipid mass spectra collected from an organoid section. Inset shows detail and complexity of the spectra (typical tissue imaging experiments result in up to thousands of such spectra). (B) Ion images generated from each peak. Each m/z value of interest is displayed as relative intensity.

**Figure S3.**
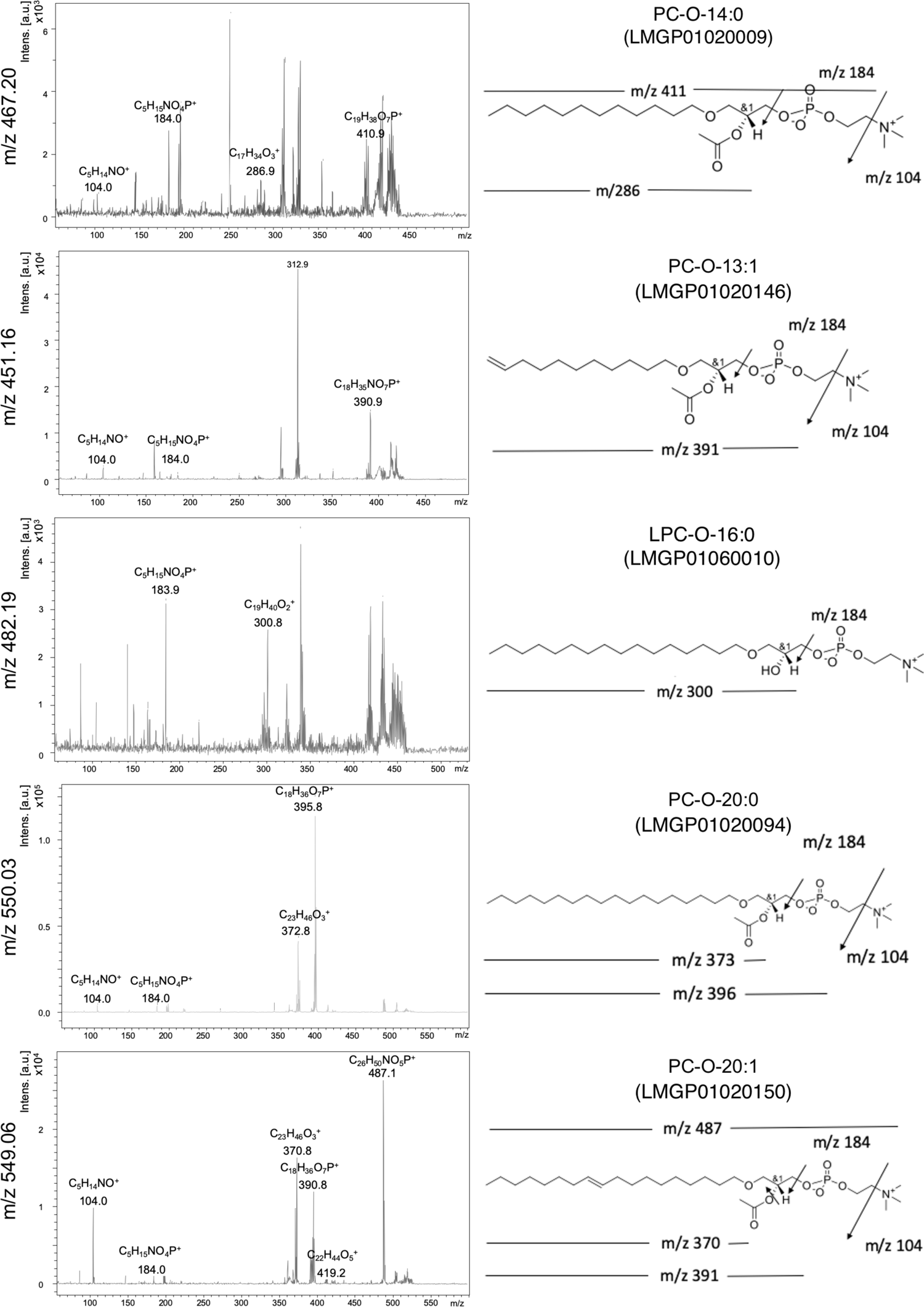
4 PTAF lipids and a PTAF lipid precursor identified by MS/MS and fragmentation analysis.

**Figure S4.**
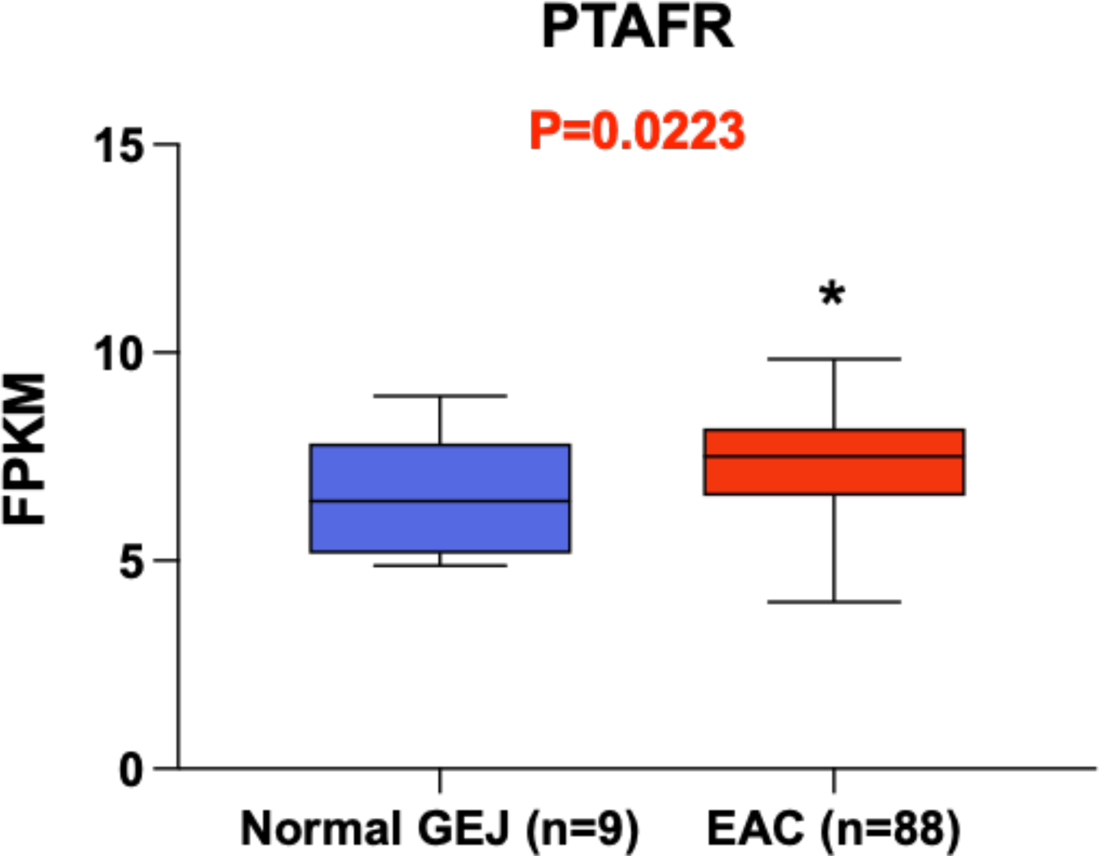
PTAFR mRNA levels in human normal GEJ *vs.* EAC tissue samples.

**Figure S5.**
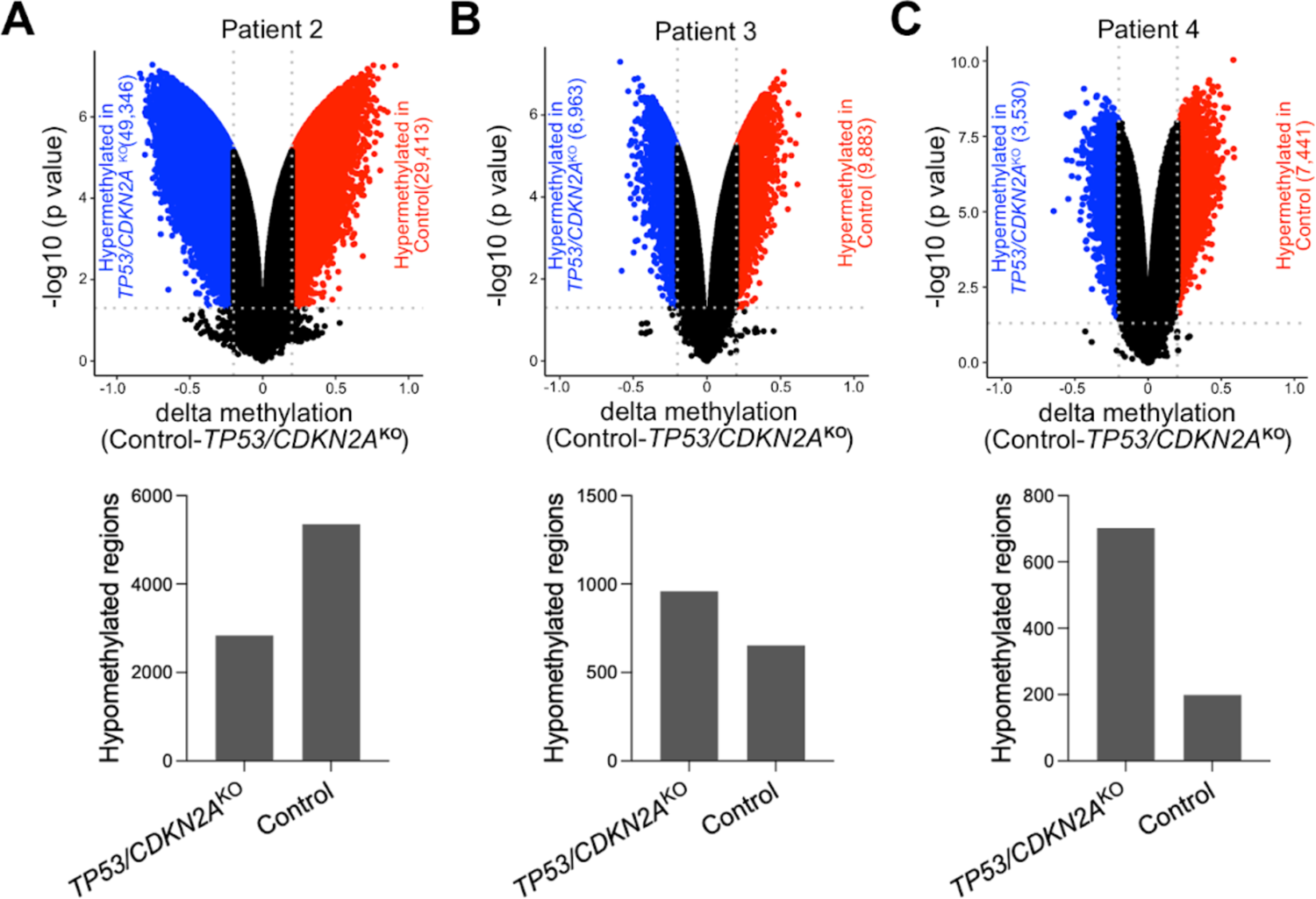
Global DNA methylation analysis in *TP53/CDKN2A*^KO^ and control GEJ organoids derived from three different patients. Volcano plots (up) revealed differentially methylated regions (DMRs), defined as exceeding a cutoff p-value of < 0.05 and an absolute delta-methylation value of > 0.2. Hypomethylated regions (*lower 3 panels*) were confirmed in *TP53/CDKN2A*^KO^ and control organoids, respectively.

